# A chemical-genetic approach for stress-independent activation of the fission yeast stress-activated protein kinase pathway

**DOI:** 10.64898/2026.06.30.735518

**Authors:** Kenneth E. Sawin, Ankita Gupta, Tatiana Dudnakova, Beste Bayrak, Adam Kovac, Domenico Modaffari, Ana I. Rodriguez-Rodriguez, Monique L. Scott, Ye Dee Tay

## Abstract

**Background:** The fission yeast stress-activated protein kinase (SAPK) pathway includes a conserved mitogen-activated protein (MAP) kinase cascade that regulates multiple cellular processes and is activated by several types of external stress. Understanding how Sty1, the MAP kinase in the SAPK pathway, controls these processes is complicated by the fact that different stressors can have stressor-specific effects that may be difficult to separate from the effects of Sty1 activation itself. Moreover, upon stress, Sty1 activation is usually short-lived. Previously, we developed a fission yeast strain, *SISA*, in which Sty1 kinase activity can be switched on in a sustained manner in the absence of external stress. This required combining multiple mutations in the SAPK pathway, including an analog-sensitive version of Sty1. When *SISA* cells are grown in the presence of analog-sensitive kinase inhibitors, Sty1 is inhibited, but when inhibitor is removed, Sty1 becomes hyperactive. While this strain was useful, it had several limitations.

**Results:** Here we describe and validate a more rationally-designed strain, *SISA4*, that retains the features of the original *SISA* strain while overcoming its limitations. *SISA4* is more stable genetically than *SISA*, easier to use in genetic crosses, and easy to identify by phenotype or genotyping. We show that analog-sensitive kinase inhibitors 4-Amino-1-tert-butyl-3-(1’-naphthylmethyl)pyrazolo[3,4-d]pyrimidine (1-NM-PP1) and 4-Amino-1-tert-butyl-3-(3-bromobenzyl)pyrazolo[3,4-d]pyrimidine (3-BrB-PP1) are equally potent for inhibiting analog-sensitive Sty1 *in vivo*, and we determine optimal inhibitor concentrations for converting *SISA4* cells from a Sty1-inhibited state to a Sty1-hyperactive state. We also find that both 1-NM-PP1 and 3-BrB-PP1 have measurable off-target effects in wild-type cells, although these are modest and generally do not affect interpretation of experiments. Finally, using SISA4, we show that the Sty1-activated transcription factor Atf1 plays an unexpected role in maintaining cell-polarity disruption after Sty1 hyperactivation.

**Conclusions:** *SISA4* will be useful for investigating how SAPK pathway activation regulates diverse cellular processes.

## INTRODUCTION

Understanding how cells respond and adapt to external stresses is fundamental to eukaryotic cell biology. In fission yeast *Schizosaccharomyces pombe*, the stress-activated protein kinase (SAPK) pathway is critical for cell adaptation and survival upon exposure to a wide variety of external stressors, including hyperosmotic and oxidative stress, heat/cold shock, nutrient deprivation, and certain heavy metals and metalloids **(Guo *et al*, 2016; Perez & Cansado, 2010; Perez *et al*, 2020; Soto *et al*, 2002)**. The SAPK pathway includes a mitogen-activated protein kinase (MAPK) cascade that is conserved from yeast to humans **(Brewster & Gustin, 2014; Canovas & Nebreda, 2021; Shabardina *et al*, 2023)**. In fission yeast, the downstream MAPK in this cascade is Sty1 (also known as Spc1/Phh1; **(Kato *et al*, 1996; Millar *et al*, 1995; Shiozaki & Russell, 1995a)**); Sty1 homologs in budding yeast and human are Hog1 and p38, respectively **(Brewster & Gustin, 2014; Canovas & Nebreda, 2021; de Nadal & Posas, 2022; Han *et al*, 2020; Martinez-Limon *et al*, 2020; Saito & Posas, 2012)**. Upon exposure to the relevant stressors, Sty1 is activated via phosphorylation of residues T171 and Y173 within its activation loop, by MAP kinase kinase (MAPKK) Wis1 **(Gaits *et al*, 1998; Millar *et al*., 1995; Samejima *et al*, 1998; Shieh *et al*, 1998; Shiozaki & Russell, 1995a; Warbrick & Fantes, 1991)**. Wis1 itself is activated via phosphorylation of residues S469 and T473 within its activation loop, by MAP kinase kinase kinases (MAPKKKs) Wis4 (also known as Wak1/Wik1) and Win1 **(Samejima *et al*, 1997; Samejima *et al*., 1998; Shieh *et al*., 1998)**. Once activated, Sty1 interacts with and phosphorylates a range of target proteins to coordinate changes in gene expression—mainly transcriptional, but also post-transcriptional—and in cell-cycle progression, to allow adaptation to stress (core environmental stress response; CESR) **(Asp *et al*, 2008; Berlanga *et al*, 2010; Chen *et al*, 2003; Gaits *et al*., 1998; Gao *et al*, 2013; Lopez-Aviles *et al*, 2005; Lopez-Aviles *et al*, 2008; Perez & Cansado, 2010; Perez *et al*., 2020; Prieto-Ruiz *et al*, 2020; Rodriguez-Gabriel *et al*, 2003; Rubio *et al*, 2021; Salat-Canela *et al*, 2017; Shiozaki & Russell, 1996; Wilkinson *et al*, 1996)**. Importantly, these changes are transient, so that once cells have adapted to a given stress (which itself is often transient), they can return to a normal state of growth and division. The transient nature of Sty1 activation depends in part on negative feedback involving dephosphorylation of Sty1’s activation loop by threonine phosphatases Ptc1 and Ptc3 **(Nguyen & Shiozaki, 1999; Shiozaki & Russell, 1995b)**, tyrosine phosphatase Pyp1 **(Millar *et al*., 1995)**, and, in particular, tyrosine phosphatase Pyp2 **(Millar *et al*., 1995)**, whose expression is highly upregulated as a result of Sty1 activation **(Chen *et al*., 2003; Degols *et al*, 1996; Rubio *et al*., 2021; Shiozaki & Russell, 1996; Wilkinson *et al*., 1996)**.

In previous work we showed that in addition to controlling stress-dependent gene expression and cell-cycle progression, the fission yeast SAPK pathway also regulates cell polarity **(Mutavchiev *et al*, 2016)**. This was demonstrated by imaging CRIB-3mCitrine, a fluorescent reporter for the active (GTP-bound) form of the Rho-family GTPase Cdc42, a key cell-polarity regulator in fission yeast **(Chiou *et al*, 2017; Martin & Arkowitz, 2014)**. Fission yeast cells are cylindrical and grow at their tips in a polarized manner. In our previous experiments, we showed that treatment with the actin-depolymerizing drug latrunculin A (LatA) leads to disruption of cell polarity, as evidenced by dispersal of CRIB-3mCitrine from the plasma membrane at cell tips and the appearance of CRIB-3mCitrine patches on the membrane at cell sides **(Mutavchiev *et al*., 2016);** see also **(Bendezu & Martin, 2011; Bendezu *et al*, 2015; Tatebe *et al*, 2008)**. We further showed that LatA treatment also leads to activation of Sty1 and that in *sty1Δ* cells, LatA treatment does *not* lead to disruption of cell polarity. Collectively, these results suggested that cell-polarity disruption after LatA treatment is driven by Sty1 activation rather than by actin depolymerization itself. Accordingly, in our previous work we also showed that Sty1 activation alone is sufficient to disrupt cell polarity. For this purpose, we developed a strain called *SISA* (for “Stress-Independent Sty1 Activation”), in which Sty1 can be activated in the absence of any external stressor **(Mutavchiev *et al*., 2016)**. The *SISA* strain brought together four distinct mutations in the SAPK pathway: *sty1-T97A*, *wis1-DD*, *pyp1Δ*, and *pyp2Δ*. Sty1-T97A (also called Sty1-as2) is an “analog-sensitive kinase” (AS-kinase) mutant of Sty1 **(Zuin *et al*, 2010)** that contains a T97A mutation within the ATP-binding pocket, allowing binding of bulky ATP-competitive adenine analogs such as 1-NM-PP1 (4-Amino-1-tert-butyl-3-(1’-naphthylmethyl)pyrazolo[3,4-d]pyrimidine) **(Bishop *et al*, 2000)**. Such analogs (here referred to as AS-kinase inhibitors) selectively inhibit AS-kinases *in vitro* with *K*_i_ values in the nanomolar range; they are also membrane permeable and thus widely used to inhibit AS-kinase activity *in vivo* **(Bishop *et al*., 2000; Cipak *et al*, 2011; Lopez *et al*, 2014; Zhang *et al*, 2013)**. Wis1-DD is a phosphomimetic mutant of Wis1 that contains mutations S469D and T473D in its activation loop, rendering it active even under non-stress conditions; consequently, *wis1-DD* cells have increased basal levels of Sty1 activation-loop phosphorylation compared to wild-type (*wis1*+) cells **(Shiozaki *et al*, 1998)**. Pyp1 and Pyp2, as mentioned above, dephosphorylate Sty1 activation-loop residue Y173, facilitating recovery from SAPK activation **(Millar *et al*., 1995; Nguyen & Shiozaki, 1999; Shiozaki & Russell, 1995a)**. By combining *sty1-T97A*, *wis1-DD*, *pyp1Δ*, and *pyp2Δ* mutations into a single strain (*SISA*) and growing it in the presence of an AS-kinase inhibitor, we were able to generate cells in which Sty1 could be phosphorylated under non-stress conditions, and thus “poised” to be active, but nevertheless inhibited in its kinase activity. Conversely, by removing the AS-kinase inhibitor, Sty1 could be rapidly switched to an active state that would persist for long periods (here referred to as “Sty1 hyperactivation”), due to the absence of negative feedback normally provided by Pyp1 and Pyp2. Using this system, we showed that upon hyperactivation of Sty1 in the *SISA* strain, CRIB-3mCitrine dispersed from cell tips and appeared in patches on cell sides, just as was observed after LatA treatment **(Mutavchiev *et al*., 2016)**.

Although the *SISA* strain makes it possible to separate the specific effects of Sty1 kinase activation from any side effects associated with individual stressors, it also presents some challenges as a tool for general use in studying the SAPK pathway. Three of the four *SISA* mutations were originally constructed in the 1990s and are tightly linked to selectable markers (two to *ura4+*, and one to *LEU2*). This makes it difficult to follow the mutations in genetic crosses, especially as the original *SISA* strain contains the CRIB-3mCitrine fluorescent reporter gene, also linked to *LEU2*. Furthermore, the *pyp1* and *pyp2* mutants are gene disruptions rather than gene deletions, and the mutant *wis1-DD* gene contains additional flanking *wis1* sequences that were later found to affect genotype/phenotype stability (see Results, below). In addition, for reasons that were initially unclear, in crosses of the *SISA* strain with other strains, progeny containing all four *SISA* mutations were recovered at frequencies much lower than expected (see Results, below). Thus, in spite of its value in helping to define the specific consequences of Sty1 activation within the SAPK pathway, improvements to the *SISA* strain would be desirable.

Here, we construct and validate a new, rationally designed, markerless fission yeast strain, called “*SISA4*”, that retains the key features of the original *SISA* strain but without any of the disadvantages mentioned above. Using the *SISA4* strain, we show that AS-kinase inhibitors 3-BrB-PP1 and 1-NM-PP1 are equipotent for inhibition of Sty1 *in vivo*, and we determine optimal conditions for their use in experiments to inhibit and hyperactivate Sty1. Surprisingly, we also find that both 3-BrB-PP1 and 1-NM-PP1 have small but measurable off-target effects in fission yeast, although these effects are sufficiently minor that they usually can be addressed through appropriate control experiments. In addition, we solve the problem associated with crossing *SISA* strains with other strains, and we provide a detailed protocol for crossing.

We also used the *SISA4* strain to investigate whether stress-induced gene expression is required for Sty1 hyperactivation-induced disruption of cell polarity, which was not previously studied with the original *SISA* strain. We find that cell-polarity disruption after Sty1 hyperactivation does not require new protein synthesis. Consistent with this, we show that the bZIP transcription factor Atf1, a key target of Sty1 **(Shiozaki & Russell, 1996; Wahls & Smith, 1994; Wilkinson *et al*., 1996)**, is not required for initial cell-polarity disruption after Sty1 hyperactivation. Surprisingly, however, we also find that longer-term, sustained cell-polarity disruption after Sty1 hyperactivation does require Atf1, although it does not require Atf1’s heterodimeric binding partner, Pcr1 **(Kanoh *et al*, 1996; Kon *et al*, 1997; Wahls & Smith, 1994; Watanabe & Yamamoto, 1996)**. While the specific function of Atf1 in cell-polarity disruption remains unclear, these results highlight the complex roles of Atf1 and Pcr1 in mediating outputs of the SAPK pathway **(Bandyopadhyay *et al*, 2017; Gaits *et al*., 1998; Gao *et al*., 2013; Lawrence *et al*, 2007; Sanso *et al*, 2008)**.

## RESULTS

### SISA4 and SISA3 strain construction

To construct a markerless *SISA* strain (**Figure 1A, B**), we used CRISPR-Cas9 **(Torres-Garcia *et al*, 2020)** to delete coding sequences for tyrosine-phosphatases Pyp1 and Pyp2 in a *sty1-T97A* mutant **(Zuin *et al*., 2010)**. In parallel, we excised *ura4+* and flanking sequences from a *wis1DD-12myc:ura4+* strain **(Shiozaki *et al*., 1998)** to generate a markerless, untagged *wis1-DD* strain (**Extended data Figure S1A**), and we crossed this with *sty1-T97A.* We then crossed the *sty1-T97A wis1-DD* strain with *sty1-T97A pyp1Δ pyp2Δ*. After tetrad dissection, spores were germinated on YE5S plates containing AS-kinase inhibitor 3-BrB-PP1, and then the resulting colonies were replica-plated onto plates lacking inhibitor.

**Figure 1.**
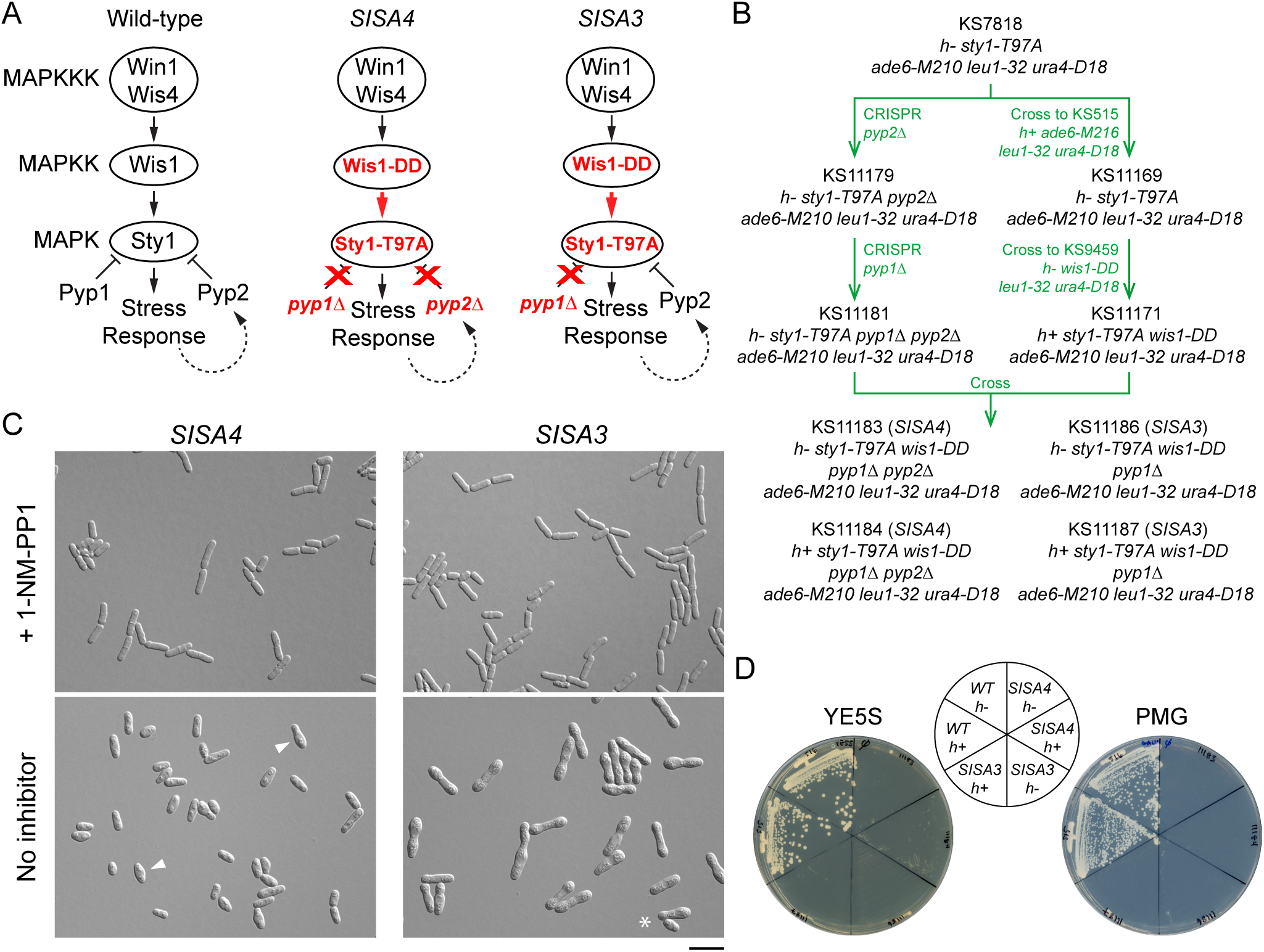
SISA4 and SISA3 strain construction and phenotype. **A.** Schematic of SAPK pathway mutations in *SISA4* and *SISA3* strains. Mutations are indicated in red. Wis1-DD is constitutively active. Sty1-T97A is inhibitable by AS-kinase inhibitors. Tyrosine phosphatases Pyp1 and Pyp2 inactivate Sty1. Pyp2 expression is positively regulated by Sty1 as part of stress response. See main text for details. **B.** Method of *SISA4* strain construction. All strains used in this work are described in **Extended data Table S2**. **C.** Morphology of *SISA4* and *SISA3* cells on YE5S agar plates in the presence of 2 µM AS-kinase inhibitor 1-NM-PP1, and after one day on plates lacking inhibitor. On plates lacking inhibitor, both strains stop dividing. *SISA4* cells become wider and swollen (arrowheads), while *SISA3* cells become larger and peanut-shaped. Some *SISA3* cells may divide at least once in the absence of inhibitor (asterisk), although they do not form microcolonies. Scale bar, 20 µm. See also **Extended data Figure S1** and **Extended data Table S1**. **D.** Absence of colony formation in *SISA4* and *SISA3* cells when streaked to rich medium (YE5S) or minimal medium (PMG) plates lacking AS-kinase inhibitor. *SISA4* cells show no growth at all. The single colony seen in the *SISA4 h-* strain is a rare revertant. Limited growth of *SISA3* cells is apparent at the primary streak site on both YE5S and PMG plates. Images of the same plates are reproduced in **Extended data Figure S2B**.

On replicas lacking inhibitor, 25 of 90 colonies (28%) showed little or no visible growth (**Extended data Figure S1B; Extended data Table S1**), indicating Sty1 hyperactivation similar to the original *SISA* strain **(Mutavchiev *et al*., 2016)**. This occurred at the expected Mendelian frequency (i.e. about 1 in 4 colonies), assuming close genetic linkage of *pyp1* and *pyp2* loci, which are only 44 kb apart (https://www.pombase.org/). However, by microscopy we found that the 25 non-growing colonies fell into two distinct classes: 18 colonies contained slightly swollen cells that were completely defective in cell division, while 7 colonies contained larger, peanut-shaped cells that appeared to retain a very limited capacity for division (**Figure 1C, D**; **Extended data Figure S1C**). Genotyping PCR revealed that this difference was due to recombination between *pyp1* and *pyp2*: all slightly swollen cells were *sty1-T97A wis1-DD pyp1Δ pyp2Δ*, while all peanut-shaped cells were *sty1-T97A wis1-DD pyp1Δ (i.e. pyp2+)* (**Extended data Table S1**). The extent of recombination between *pyp1* and *pyp2* loci was at first surprising, as only about 7% *pyp1-pyp2* recombinants were expected, assuming a genetic distance of approximately 1 centiMorgan per ∼6.6 kb in *S. pombe* **(Egel, 2004; Munz *et al*, 1989)**. However, genotyping PCR of all 90 colonies revealed ∼21% *pyp1-pyp2* recombinants, confirming the higher-than-expected recombination frequency (**Extended data Table S1**). As a previous genome-wide analysis of meiotic DNA double-strand breaks (DSBs) identified a DSB hotspot, plus two weaker hotspots, in the *pyp1-pyp2* interval **(Fowler *et al*, 2014)**, this likely accounts for the increased number of *pyp1*-*pyp2* recombinants observed.

Overall, the phenotype of the quadruple mutant *sty1-T97A wis1-DD pyp1Δ pyp2Δ* on plates lacking AS-kinase inhibitor appears identical to that of our original *SISA* strain **(Mutavchiev *et al*., 2016)**, while that of the triple mutant *sty1-T97A wis1-DD pyp1Δ* is similar but distinct. By contrast, the triple mutant *sty1-T97A wis1-DD pyp2Δ* did not have a striking phenotype (**Extended data Table S1**), consistent with the view that Pyp1 may have a generally more prominent role than Pyp2 **(Millar *et al*, 1992; Ottilie *et al*, 1992).** Based on the number of SAPK-relevant mutations in our new strains, and to distinguish them from our original *SISA* strain, we will refer to the quadruple mutant *sty1-T97A wis1-DD pyp1Δ pyp2Δ* as *SISA4* and to the triple mutant *sty1-T97A wis1-DD pyp1Δ* as *SISA3*.

### SISA4 phenotype stability

With our original *SISA* strain **(Mutavchiev *et al*., 2016)**, when cells were streaked directly onto plates lacking AS-kinase inhibitor—which is normally lethal for *SISA* cells—rare colonies could occasionally form. This phenotype reversion could result from a loss-of-function mutation in *sty1* or *wis1* or, alternatively, from a spontaneous loop-out excision of *ura4+* at the *wis1* locus, which can revert *wis1DD-12myc* to *wis1+* (**Extended data Figure S1A**). Given that *SISA4* cells already have *ura4+* excised from the *wis1* locus (**Extended data Figure S1A)**, we reasoned that if loop-out excision of *ura4+* were the major cause of phenotype reversion in the original *SISA* strain, then the *SISA4* strain should have a much lower frequency of phenotype reversion than the original *SISA* strain. We developed a quantitative assay for phenotype reversion and found that this was indeed the case (**Figure 2A, B; see Methods**). Based on a simple model, we estimate that the reversion frequency in the original *SISA* strain is approximately 5 x 10^-6^ per cell per generation, while in *SISA4* it is up to about 500 times lower (**see Methods**). This demonstrates that *SISA4* is much more stable genetically than the original *SISA* strain and that excision of *ura4+* at the *wis1* locus is indeed the major cause of reversion in the original *SISA* strain.

**Figure 2.**
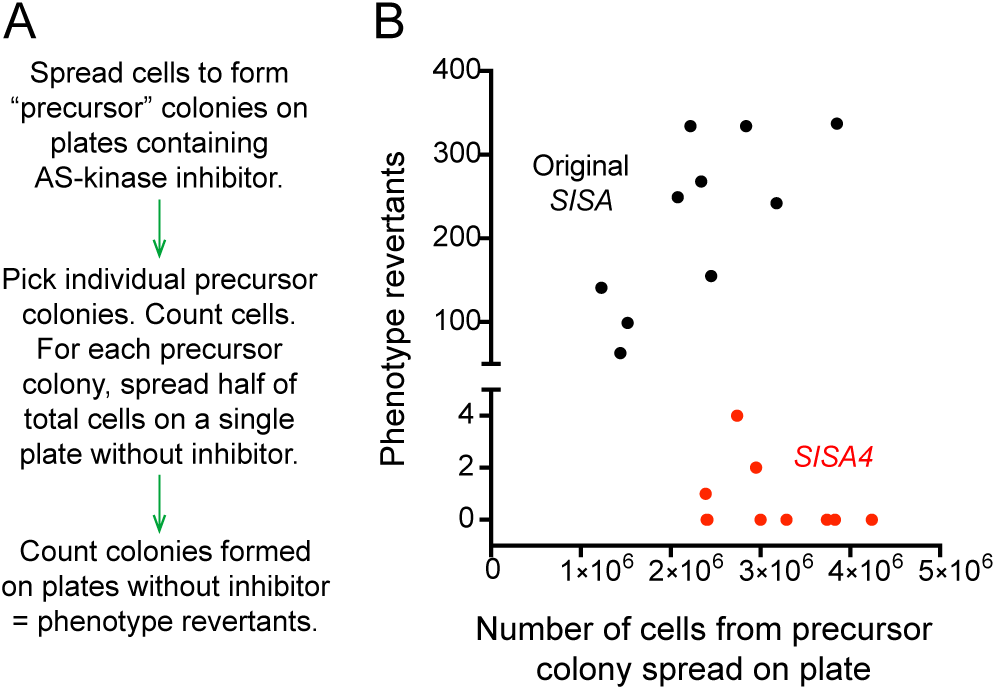
*SISA4* phenotype reversion frequency. **A.** Outline of phenotype reversion assay. **B.** Quantitation of phenotype reversion in *SISA4* cells compared to original SISA strain from Mutavchiev *et al*. (2016).

### Optimal AS-kinase inhibitor concentrations for use with SISA4 and SISA3 strains

We next investigated what concentrations of AS-kinase inhibitors are most suitable for use in *SISA4* and *SISA3* strains. Because AS-kinase inhibitors are expensive and may also have off-target effects (see below), we sought to identify the lowest inhibitor concentrations that allow robust growth of *SISA* strains without significant Sty1 activation, as a major use of *SISA* strains is to study how cells respond to acute increases in Sty1 activity upon removal of inhibitor.

First, we assayed colony size on rich and minimal medium plates containing different concentrations of AS-kinase inhibitors (**Figure 1D**; **Figure 3; Extended data Figure S2**). We tested both 3-BrB-PP1 **(Cipak *et al*., 2011; Okuzumi *et al*, 2010)**, which was used with our original *SISA* strain **(Mutavchiev *et al*., 2016)**, and 1-NM-PP1 **(Bishop *et al*., 2000)**, which is normally less expensive than 3-BrB-PP1 and more frequently used to inhibit AS-kinases. We obtained similar results on rich vs. minimal medium, and similar results with 3-BrB-PP1 vs. 1-NM-PP1. On plates lacking AS-kinase inhibitors, neither *SISA4* nor *SISA3* cells formed visible colonies. By contrast, at 0.05 µM and higher inhibitor concentrations, both *SISA4* and *SISA3* cells formed colonies. At inhibitor concentrations between 0.05 µM and 0.5 µM, *SISA4* and *SISA3* colonies were smaller than wild-type colonies, and colony size increased with inhibitor concentration. This suggests that within this concentration range, Sty1 activity in *SISA4* and *SISA3* cells is higher than normal and partially inhibitory to cell growth, and that as inhibitor concentration increases, Sty1 activity decreases. Within this concentration range, we also found that *SISA3* colonies were nearly always larger than *SISA4* colonies. This is consistent with the partial attenuation of Sty1 activity by Pyp2, which itself is induced by Sty1 activation **(Chen *et al*., 2003; Degols *et al*., 1996; Rubio *et al*., 2021; Shiozaki & Russell, 1996; Wilkinson *et al*., 1996)**; it also agrees with our observations that *SISA3* cells may exhibit some very limited growth in the absence of inhibitor, although not enough to form a visible colony (**Figure 1C, D; Extended data Figure S1C; Extended data Figure S2B**). At 1 µM and higher AS-kinase inhibitor concentrations (up to 10 µM), *SISA4* and *SISA3* colony size was essentially the same as in wild-type cells, suggesting that Sty1 is not overly active at these concentrations.

**Figure 3.**
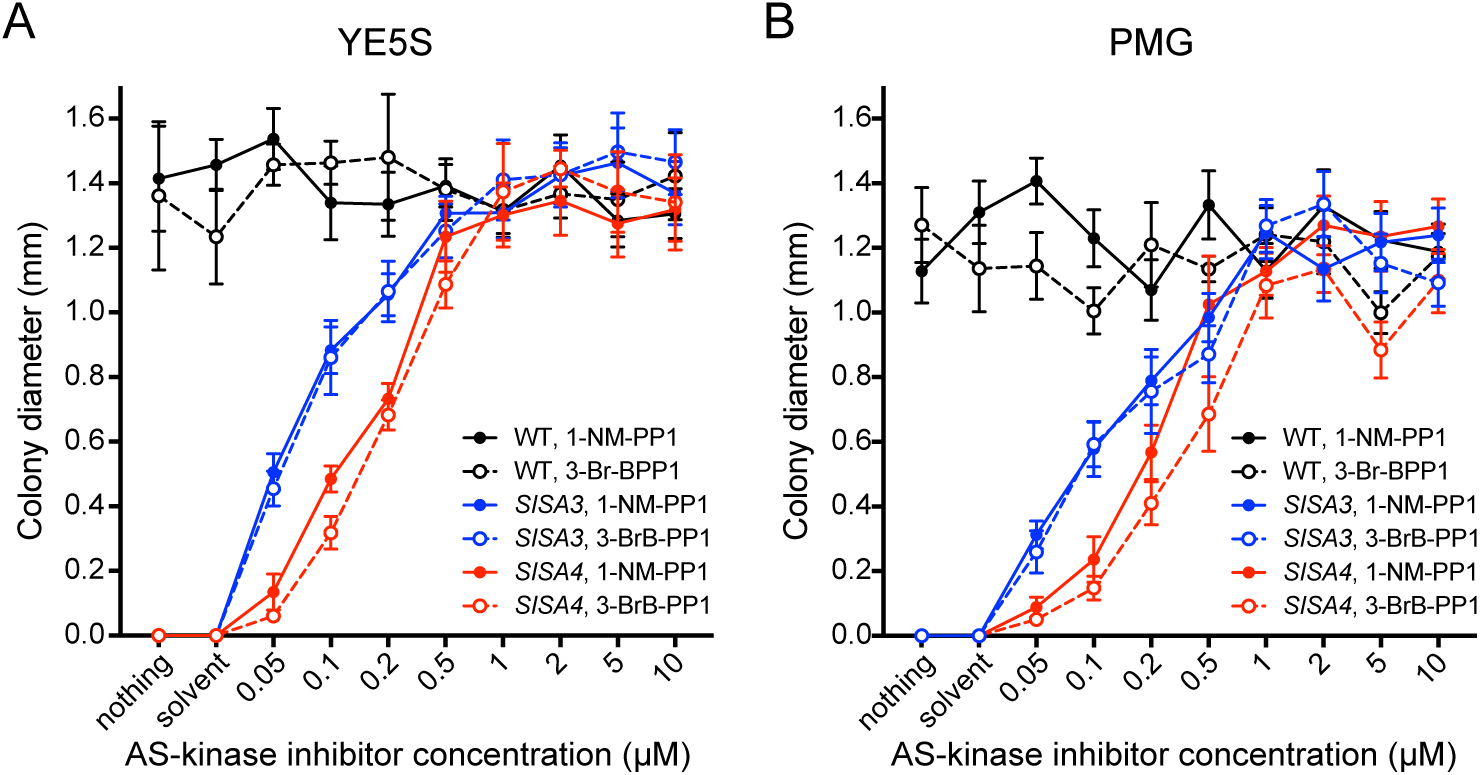
*SISA4* and *SISA3* colony size matches wild-type at 1 µM and higher AS-kinase inhibitor concentrations. Colony diameters for wild-type (WT; i.e. non-*SISA*), *SISA3*, and *SISA4* strains after 3 days (30°C) on (A) rich medium (YE5S) and (B) minimal medium (PMG) plates containing different concentrations of AS-kinase inhibitors 1-NM-PP1 and 3-BrB-PP1. Each data point shows mean for 10 colonies. Error bars indicate SD. Images of plates are shown in **Extended data Figure S2**.

Second, focusing on *SISA4*, we measured cell length at septation in wild-type, *SISA4*, and *sty1Δ* cells grown in different concentrations of AS-kinase inhibitors **(Figure 4)**. While colony-size assays suggested that Sty1 is not overly active in *SISA4* cells at 1-10 µM inhibitor concentrations, they did not address whether Sty1 is more strongly inhibited at the higher vs. the lower end of this range. Because *sty1* loss-of-function mutants divide at a greater length than wild-type cells, especially in minimal medium; **(Millar *et al*., 1995; Shiozaki & Russell, 1995a)**, we reasoned that cell length at septation could serve as a proxy for Sty1 inhibition in AS-kinase inhibitor-treated *SISA4* cells. We tested both 3-BrB-PP1 and 1-NM-PP1, in both rich and minimal medium. Although there was some variation in the data, we obtained similar results in all conditions. Neither wild-type nor *sty1Δ* cells showed a consistent relationship between inhibitor concentration and cell length at septation. By contrast, *SISA4* cells showed a general monotonic relationship between AS-kinase inhibitor concentration and cell length at septation, especially in minimal medium, and cell length at septation was approximately the same at 2 µM, 5 µM, and 10 µM inhibitor concentrations (**Figure 4**). This suggests that inhibitor concentrations greater than 10 µM would not lead to further increased length at septation, and it further suggests that at 2 µM inhibitor concentration, Sty1 (i.e. Sty1-T97A) is probably as inhibited as it can be *in vivo*. Interestingly, even at high inhibitor concentrations, cell length at septation in inhibitor-treated *SISA4* cells was generally smaller than in *sty1Δ* cells. This suggests either that some Sty1 kinase activity may remain present even at high AS-kinase inhibitor concentrations or, alternatively, that complete inhibition of Sty1 kinase activity does not produce the same septation-length phenotype as *sty1* deletion.

**Figure 4.**
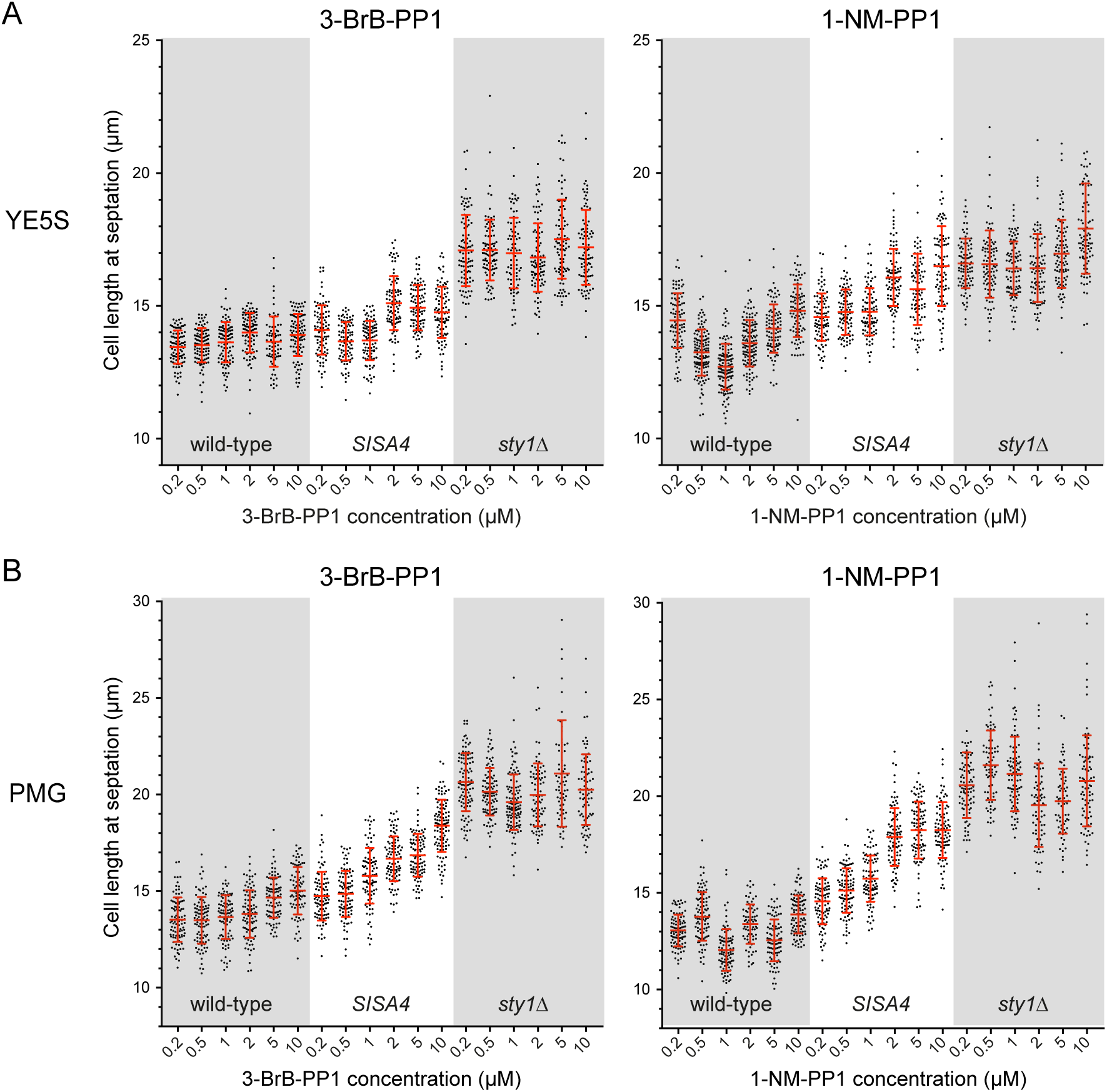
AS-kinase inhibitors 3-BrB-PP1 and 1-NM-PP1 have similar dose-dependent effects on cell length at septation in *SISA4* cells. Cell length at septation in wild-type (i.e. non-*SISA*), *SISA4*, and *sty1Δ* cells grown in (A) rich medium (YE5S) and (B) minimal medium (PMG) at the indicated concentrations of AS-kinase inhibitors 1-NM-PP1 and 3-BrB-PP1. Lines and error bars indicate mean and SD.

Third, to further assess Sty1 inhibition by AS-kinase inhibitors, we performed “inhibitor washout” experiments, in which cells were grown in the presence of AS-kinase inhibitor, harvested by filtration, and then resuspended in fresh medium containing different concentrations of inhibitor, or no inhibitor at all (**Figure 5**). As a quantitative assay for Sty1 activity, we measured fluorescence of an Lsd90-3xmNeonGreen (Lsd90-3mNG) fusion protein by flow cytometry, three hours after washout. Expression of *lsd90* is controlled by the SAPK pathway and the CESR, and *lsd90* mRNA levels are strongly increased after exposure to a variety of external stressors **(Chen *et al*., 2003; Rubio *et al*., 2021)**. We therefore predicted that after inhibitor washout in *SISA4* cells, Sty1 hyperactivation would lead to significantly increased Lsd90-3mNG expression. In a first set of experiments, we found that washout from 5 µM inhibitor (either 3-BrB-PP1 or 1-NM-PP1) to no inhibitor led to a 300- to 700-fold increase in Lsd90-3mNG fluorescence, and washout to 0.2 µM inhibitor produced a similar but slightly smaller increase (**Figure 5A**). By contrast, we did not observe any increase in Lsd90-3mNG fluorescence after mock wash (i.e. from 5 µM inhibitor to fresh medium also containing 5 µM inhibitor) or after washout from 5 µM to 2 µM inhibitor. Washout to 0.5 µM inhibitor led to an intermediate increased fluorescence, and washout to 1 µM inhibitor led to a mixed response in which some cells showed increased fluorescence while others did not. As part of a second set of experiments, we found that washout from 2 µM inhibitor (either 3-BrB-PP1 or 1-NM-PP1) to no inhibitor also led to a several hundred-fold increase in Lsd90-mNG fluorescence (**Figure 6A**).

**Figure 5.**
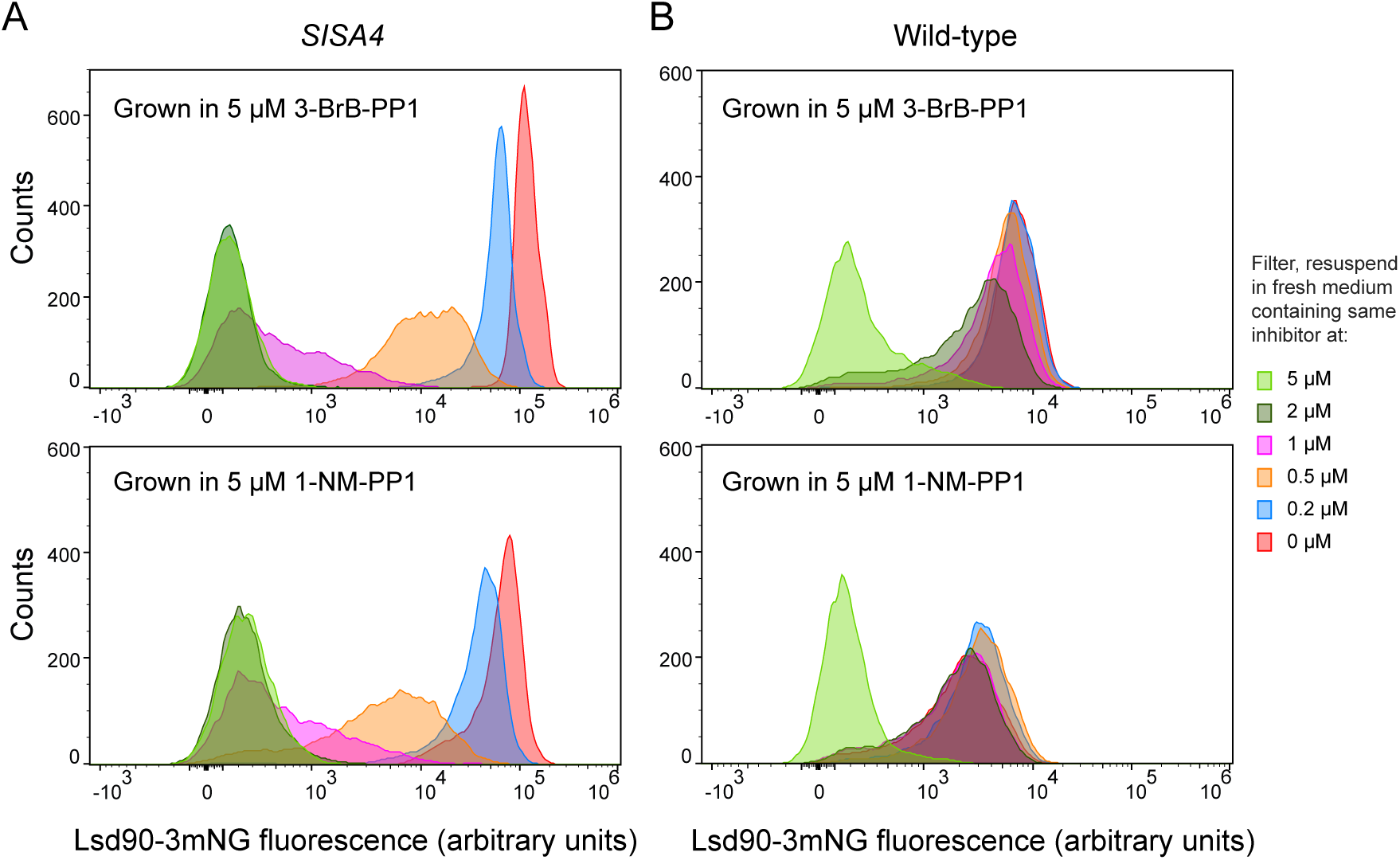
Changes in expression of core environmental stress response gene *lsd90* in *SISA4* and wild-type cells after washout of AS-kinase inhibitors. Flow cytometry analysis of Lsd90-3xmNeonGreen (Lsd90-3mNG) expression in (A) *SISA4* and (B) wild-type (i.e. non-*SISA*) cells three hours after washout from growth medium containing 5 µM 3-BrB-PP1 or 1-NM-PP1 into medium containing different concentrations of the same inhibitors, or no inhibitor (0 µM). In *SISA4* cells, Lsd90 expression increases greatly after washout to 0 or 0.2 µM inhibitor, moderately after washout to 0.5 µM, less after washout to 1 µM, and not at all after washout to 2 or 5 µM inhibitor (washout from 5 µM to 5 µM inhibitor represents a “mock wash”). This suggests that 2 µM inhibitor is as effective as 5 µM inhibitor for inhibiting Sty1-T97A kinase activity in *SISA4* cells. In wild-type cells, in all washouts except mock wash, Lsd90 expression increases, although extremely modestly compared to the increases in *SISA4* cells. This suggests that 3-BrB-PP1 and 1-NM-PP1 have mild off-target effects in wild-type cells (see main text). Key at right applies to both A and B. See also **Extended data Figures S3 and S4**.

**Figure 6.**
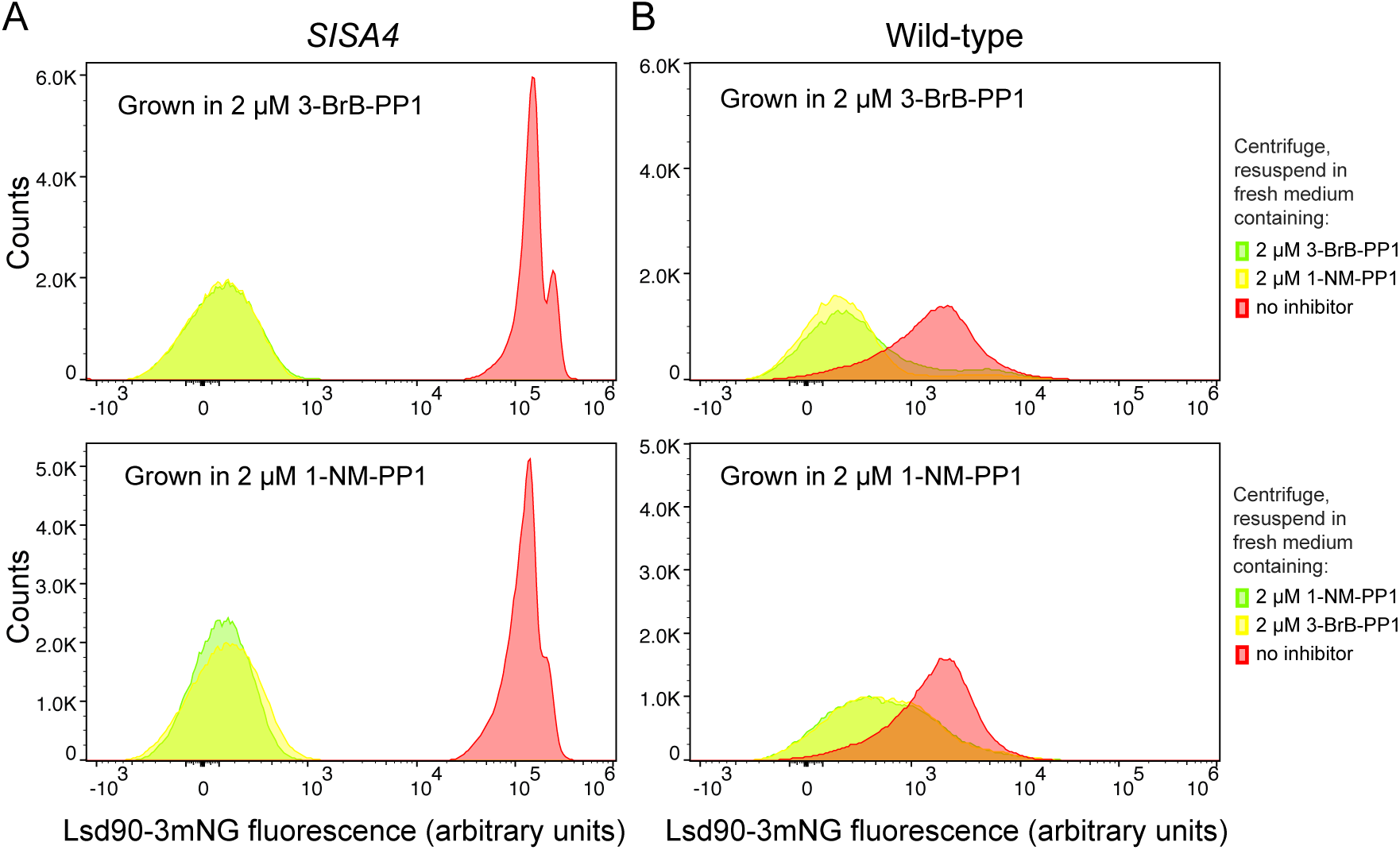
AS-kinase inhibitors 3-BrB-PP1 and 1-NM-PP1 have indistinguishable on-target effects and indistinguishable off-target effects. Flow cytometry analysis of Lsd90-3xmNeonGreen (Lsd90-3mNG) expression in (A) *SISA4* and (B) wild-type (i.e. non-*SISA*) cells three hours after “inhibitor swap”. Cells were grown in medium containing either 2 µM 3-BrB-PP1 or 2 µM 1-NM-PP1, briefly centrifuged, and then resuspended in fresh medium containing the same concentration of the same inhibitor (mock wash), the same concentration of the other inhibitor (inhibitor swap), or no inhibitor (washout). Keys at right apply to both A and B. In both *SISA4* and wild-type cells, inhibitor swap has no effect on Lsd90 expression compared to mock wash. For *SISA4* cells, this confirms that 3-BrB-PP1 and 1-NM-PP1 are equally potent for inhibition of Sty1-T97A *in vivo*. For wild-type cells, this indicates that the sensing mechanisms that lead to modest increases of Lsd90-3mNG expression after washout in wild-type cells do not distinguish between 3-BrB-PP1 and 1-NM-PP1.

Overall, our results from inhibitor washout are consistent with our analysis of cell length at septation in showing that 2 µM and 5 µM AS-kinase inhibitor concentrations have equivalent effects on Sty1 inhibition *in vivo* in *SISA4* cells. We conclude that for converting *SISA4* cells from a Sty1-inhibited state to a Sty1-hyperactivated state, washout from 2 µM to no inhibitor is likely optimal.

### AS-kinase inhibitors 3-BrB-PP1 and 1-NM-PP1 have mild but measurable off-target effects in fission yeast

Alongside AS-kinase inhibitor washout experiments in *SISA4* cells, we performed equivalent experiments in wild-type cells expressing Lsd90-3mNG (**Figure 5B**). Surprisingly, washouts from 5 µM inhibitor (either 3-BrB-PP1 or 1-NM-PP1) to no inhibitor led to a 15- to 30-fold increase in Lsd90-3mNG fluorescence. Washouts from 5 µM inhibitor to intermediate inhibitor concentrations (ranging from 0.2 to 2 µM) led to slightly smaller increases. While the increase in Lsd90-3mNG fluorescence seen in wild-type cells was modest—between one and two orders of magnitude less than the increase in *SISA4* cells—it was unexpected, and we therefore investigated it further. For technical reasons, in our next experiments, washout was performed by brief centrifugation rather than filtration, and therefore we also confirmed that centrifugation itself did not lead to increased Lsd90-3mNG expression (**Extended data Figure S3A, B**). As with filtration washout, centrifugation washout from either 5 µM or 2 µM inhibitor (3-BrB-PP1 or 1-NM-PP1) to no inhibitor or intermediate inhibitor concentrations (e.g. 0.2 µM, 0.5 µM) also led to increased Lsd90-3mNG fluorescence (**Extended data Figure S3A, B**).

Given that wild-type cells do not contain an AS-kinase, we next investigated what might be responsible for their increased Lsd90-3mNG fluorescence after AS-kinase inhibitor washout. Because *lsd90* expression increases after exposure to a variety of external stressors that activate the SAPK pathway **(Chen *et al*., 2003; Rubio *et al*., 2021),** we hypothesized that when wild-type cells are grown for long periods in the presence of AS-kinase inhibitor and then inhibitor is removed, this may also mildly activate the SAPK pathway (see Discussion). In this scenario, increased Lsd90-mNG fluorescence after washout in wild-type cells would be expected to depend on Sty1. Accordingly, we found that in *sty1Δ* cells expressing Lsd90-mNG, washout from 2 µM inhibitor (either 3-BrB-PP1 or 1-NM-PP1) to no inhibitor did not lead to any increase in Lsd90-mNG fluorescence (**Extended data Figure S3C**). We also considered an alternative hypothesis, namely that AS-kinase inhibitors themselves could inhibit wild-type Sty1, or some other protein in the SAPK/CESR pathway, such that when wild-type cells are grown in AS-kinase inhibitor (as in our washout experiments), steady-state *lsd90* expression is lower than normal basal levels, and then when inhibitor is removed, *lsd90* expression is restored to normal levels. However, Lsd90-3mNG fluorescence of wild-type cells grown in different steady-state concentrations of AS-kinase inhibitors was identical in the presence and absence of inhibitors, indicating that AS-kinase inhibitors do not inhibit the wild-type SAPK/CESR pathway (**Extended data Figure S4**).

Collectively, our AS-kinase inhibitor washout experiments in wild-type cells suggest that both 3-BrB-PP1 or 1-NM-PP1—even when used at low micromolar concentrations—can have mild but measurable off-target effects in fission yeast. However, given the extreme sensitivity of the Lsd90-3mNG reporter, and the scale of the different effects of inhibitor washout in *SISA4* vs wild-type cells **(Figure 5**; **Figure 6)**, these mild effects are unlikely to significantly affect the interpretation of inhibitor washout experiments involving Sty1 hyperactivation in *SISA4* cells (see below and Discussion).

### AS-kinase inhibitors 3-BrB-PP1 and 1-NM-PP1 have equivalent effects in S. pombe

While the interactors of 3-BrB-PP1 and 1-NM-PP1 that lead to the off-target effects described above are unknown, we used an “inhibitor swap” approach to determine whether off-target interactors are likely to be the same for the two inhibitors. We used centrifugation washout to replace 2 µM 3-BrB-PP1 with 2 µM 1-NM-PP1, and vice versa, in *SISA4* and wild-type cells expressing Lsd90-3mNG. In all cases, washout from one inhibitor to the other inhibitor did not alter Lsd90-mNG fluorescence (**Figure 6**). By contrast, washout from either inhibitor to no inhibitor led to increased Lsd90-mNG fluorescence as described above. These results suggest that the off-target interactors of 3-BrB-PP1 and 1-NM-PP1 may be identical; at minimum, at least some off-target interactors are likely to be common to the two inhibitors. Overall, our analyses in *SISA4* and wild-type cells suggest that 3-BrB-PP1 and 1-NM-PP1 are essentially interchangeable for use in *SISA4* cells. Based on cost, availability, and wide usage of 1-NM-PP1, we chose washout of 2 µM 1-NM-PP1 to no inhibitor as a standard method for converting *SISA4* cells from a Sty1-inhibited state to a Sty1-hyperactivated state.

### Using SISA4 to study dynamics of the Cdc42 cell-polarity module

We next tested how Sty1 hyperactivation in *SISA4* cells affects the spatial distribution of the Cdc42 cell-polarity module *in vivo*, using CRIB-3mCitrine to monitor localization of active (GTP-bound) Cdc42 **(Bendezu & Martin, 2011; Mutavchiev *et al*., 2016; Tatebe *et al*., 2008)**. In the original *SISA* strain, washout from 5 µM 3-BrB-PP1 to no inhibitor led to a strong decrease in CRIB-3mCitrine signal at its normal position on the plasma membrane at cell tips, as well as the appearance of dynamic patches of CRIB-3mCitrine on the plasma membrane at cell sides, accompanied by arrest of cell elongation and of cell-cycle progression **(Mutavchiev *et al*., 2016)**. To perform equivalent experiments in *SISA4 CRIB-3mCitrine* cells, we used washout from 2 µM 1-NM-PP1 to no inhibitor. Consistent with expectation, this led to the same changes in CRIB-3mCitrine localization observed after 3-BrB-PP1 washout in the original *SISA* strain (**Figure 7A**). Importantly, washout of inhibitor could be achieved with fewer washes than in our original experiments (see Methods). In control mock wash experiments (i.e. replacing medium containing 2 µM 1-NM-PP1 with fresh medium also containing 2 µM 1-NM-PP1), CRIB-3mCitrine remained at cell tips, and cells continued to elongate (**Figure 7B**).

**Figure 7.**
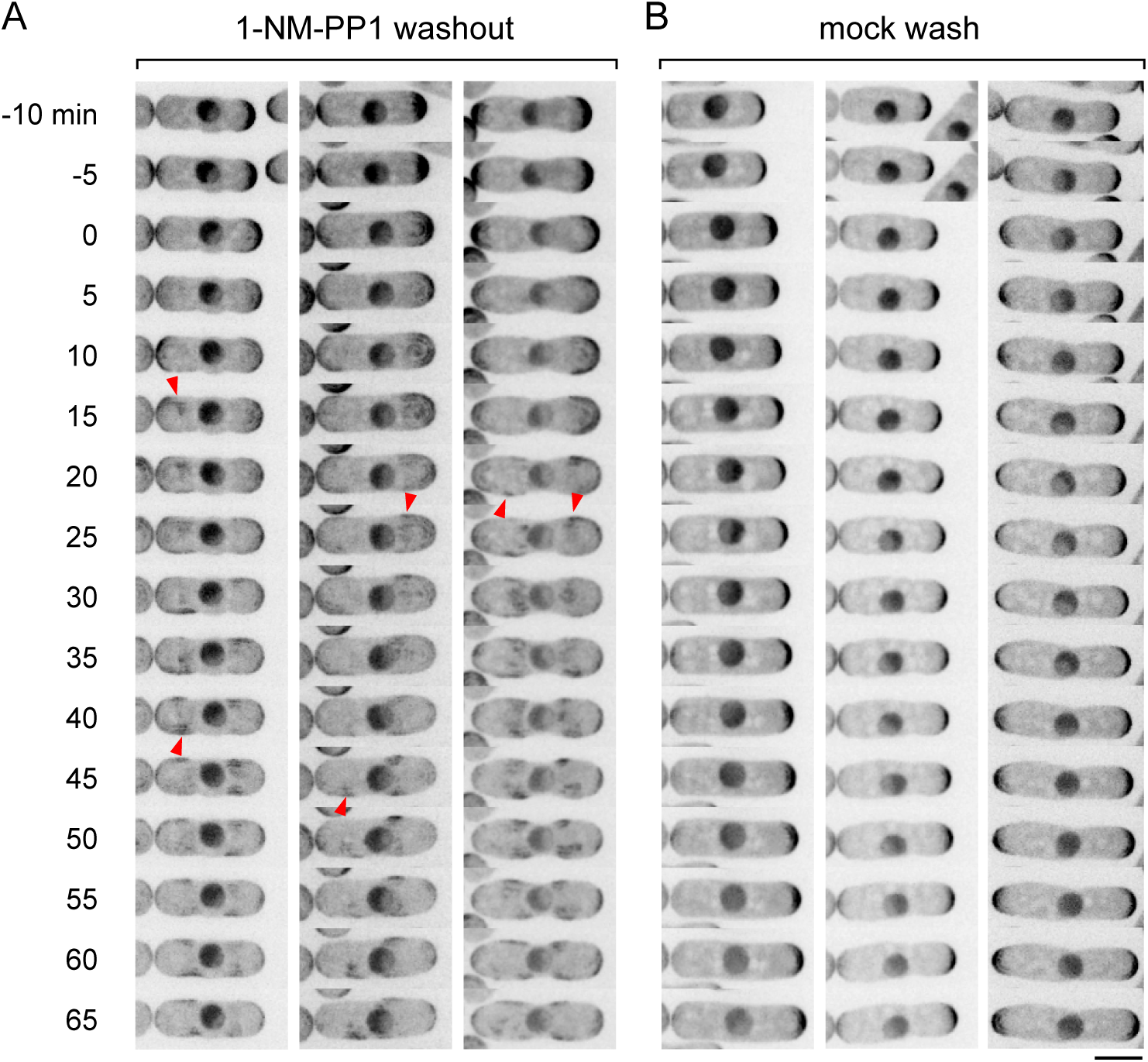
Cell polarity disruption after Sty1 hyperactivation in *SISA4* cells. Timepoints from movies of *SISA4* cells expressing Cdc42-GTP reporter CRIB-3mCitrine, after (A) washout of 2 µM 1-NM-PP1 from growth medium or (B) mock wash into growth medium still containing 2 µM 1-NM-PP1. Times are relative to washout or mock wash; because of multiple washes, the zero timepoint was defined as the first timepoint after washing (see Methods). After washout, CRIB-3mCitrine decreases at cell tips and appears in patches on cell sides (arrowheads), and cells do not elongate. Scale bar, 5 µm. See also **Extended data Figure S5**.

We also imaged CRIB-3mCitrine in wild-type cells after washout from 2 µM 1-NM-PP1 to no inhibitor. In these cells, CRIB-3mCitrine remained at cell tips, and cells continued to elongate (**Extended data Figure S5**). This indicates that even if mild activation of Sty1 did occur in wild-type cells after inhibitor washout, this is insufficient to alter the Cdc42 cell-polarity module. Overall, we conclude that changes in cell polarity seen after 1-NM-PP1 washout in *SISA4* cells are unlikely to be significantly affected by off-target effects.

Although a major role of Sty1 and the SAPK pathway is to regulate stress-dependent gene expression **(Chen *et al*., 2003; Rubio *et al*., 2021; Shiozaki & Russell, 1996; Wilkinson *et al*., 1996),** our previous work using latrunculin A as a SAPK pathway stressor suggested that Sty1 activation-induced changes in the Cdc42 cell-polarity module occur post-translationally, independent of new gene expression **(Mutavchiev *et al*., 2016)**. However, this was never tested in *SISA* cells, which offer a more precise way to manipulate Sty1 activity. We therefore imaged CRIB-3mCitrine after 1-NM-PP1 washout in *SISA4* cells that had been pre-treated with cycloheximide (CHX) to inhibit protein synthesis (**Figure 8**). After 1-NM-PP1 washout in *SISA4* CHX-treated cells, we observed strongly decreased CRIB-3mCitrine at cell tips and dynamic patches of CRIB-3mCitrine at cell sides (**Figure 8A**), i.e. the same changes that are seen after washout in non-CHX-treated cells. These changes were not due to CHX treatment itself, because after mock wash in *SISA4* CHX-treated cells, CRIB-3mCitrine remained at cell tips, although cell elongation was largely inhibited, presumably because of inhibition of protein synthesis (**Figure 8B**). Overall, these results indicate that new protein synthesis is not required for the changes in the spatial distribution of the Cdc42 cell-polarity module seen after Sty1 hyperactivation in SISA4 cells, consistent with our previous results using CHX in LatA-treated cells **(Mutavchiev *et al*., 2016)**.

**Figure 8.**
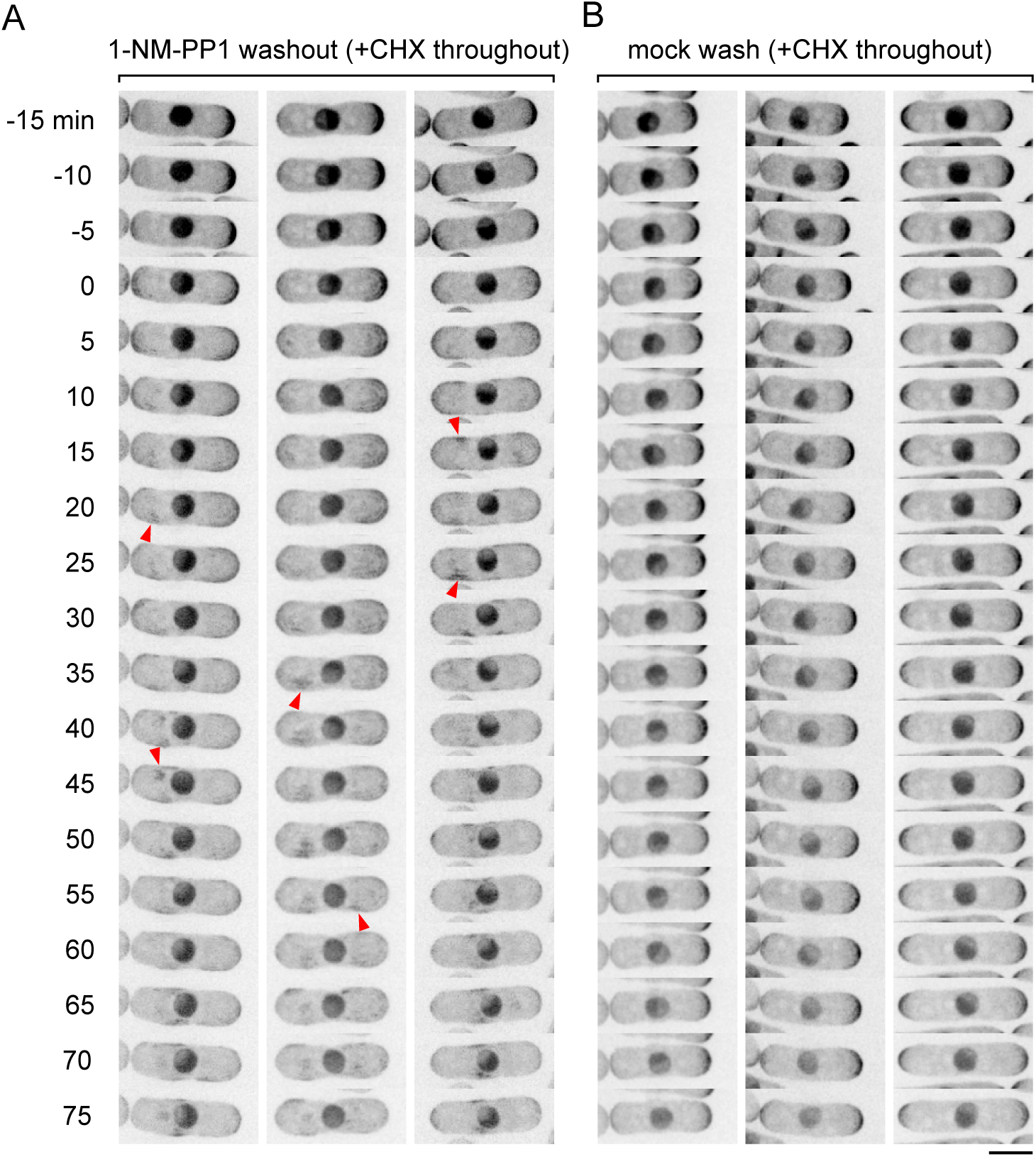
Sty1 hyperactivation-induced cell polarity disruption does not require new protein synthesis. Timepoints from movies of cycloheximide (CHX)-treated *SISA4* cells expressing CRIB-3mCitrine, after (A) washout of 2 µM 1-NM-PP1 from growth medium or (B) mock wash into growth medium still containing 2 µM 1-NM-PP1. CHX was present both before and after washout and mock wash (see Methods). Times are as in Figure 7. Arrowheads show examples of CRIB-3mCitrine patches on cell sides. CHX treatment alone (i.e. in mock-washed cells) does not affect CRIB-3mCitrine localization. Scale bar, 5 µm.

We also imaged CRIB-3mCitrine after 1-NM-PP1 washout in *SISA4 atf1Δ* cells. Atf1 is phosphorylated by Sty1 upon SAPK pathway activation and is required for most SAPK pathway-dependent changes in gene expression **(Chen *et al*., 2003; Sanso *et al*., 2008).** We previously showed that after LatA treatment of *atf1Δ* cells, changes in CRIB-3mCitrine localization occur as they do in wild-type cells **(Mutavchiev *et al*., 2016)**. Consistent with this, in *SISA4 atf1Δ* cells, 1-NM-PP1 washout initially led to changes in CRIB-3mCitrine localization similar to those seen after washout in *SISA4* (*atf1+*) cells (**Figure 9, Extended data Figure S6**). This indicates that Atf1 is not required for Sty1 hyperactivation-induced changes in the spatial distribution of the Cdc42 cell-polarity module. However, at later imaging times/timepoints, we observed at least partial recovery of CRIB-3mCitrine to cell tips in approximately 30% of *SISA4 atf1Δ* cells (**16 of 55 cells; Figure 9)**, and after Sty1 hyperactivation for one day, essentially all *SISA4 atf1Δ* cells were highly elongated compared to *SISA4* (*atf1+*) cells, indicating some recovery of polarized growth (**Figure 10E, F**). After an additional day of Sty1 hyperactivation, *SISA4 atf1Δ* cells did not elongate further (**Figure 10F, J**), although they swelled in the cell central region; this may be analogous to the extensive swelling seen in *SISA4* (*atf1+*) cells after two days of Sty1 hyperactivation (**Figure 10I**). Interestingly, in spite of recovery of polarized growth, *SISA4 atf1Δ* cells showed no signs of cell division, even after two days of Sty1 hyperactivation, suggesting that Sty1 was still sufficiently hyperactive to prevent cell-cycle progression (**Figure 10B, F, J**).

**Figure 9.**
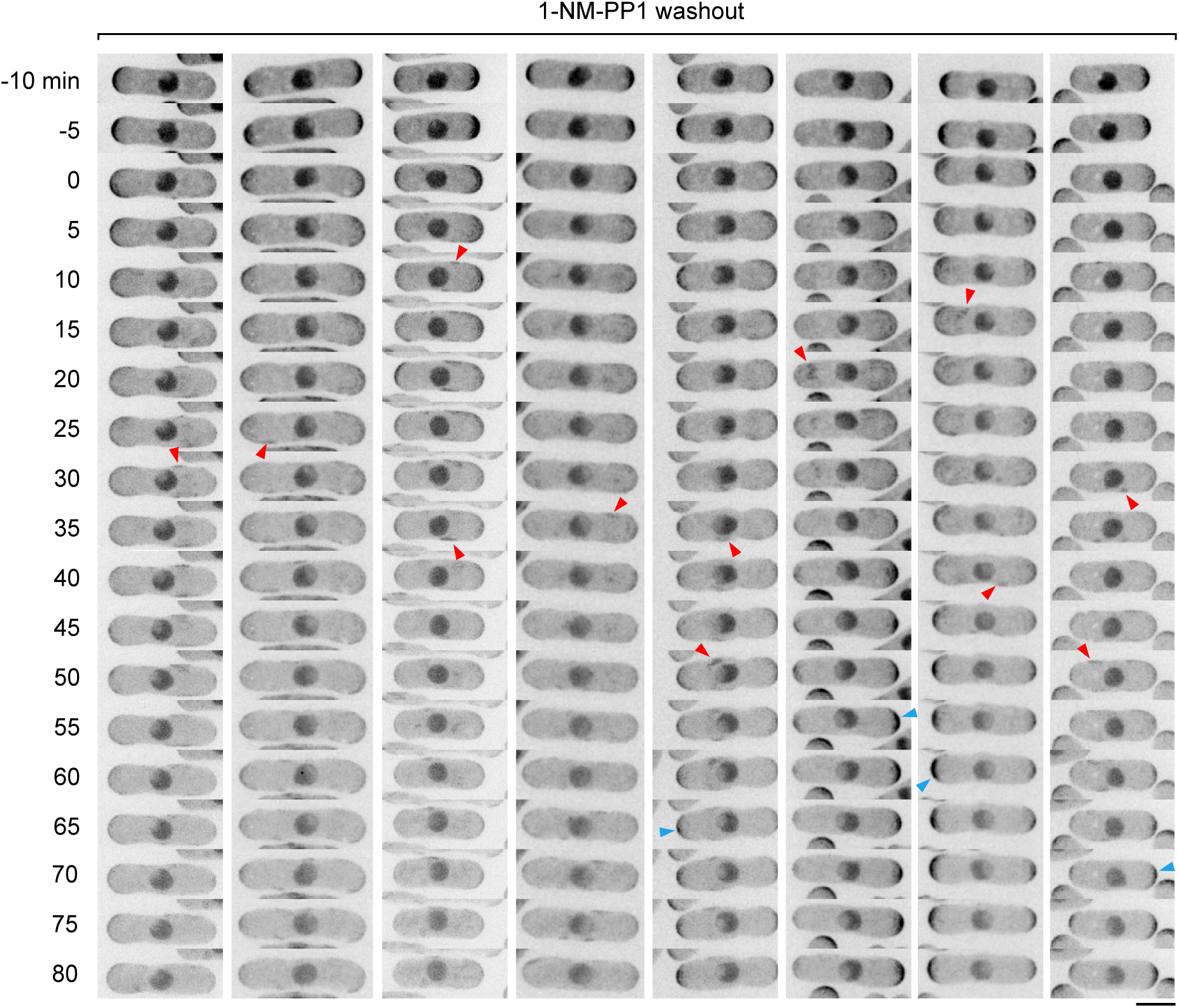
Deletion of stress-activated transcription factor Atf1 allows partial recovery of Cdc42-GTP to cell tips in *SISA4* cells after Sty1 hyperactivation-induced polarity disruption. Timepoints from movies of *SISA4 atf1Δ* cells expressing CRIB-3mCitrine, after washout of 2 µM 1-NM-PP1 from growth medium. Times are as in Figures 7 and 8. After washout, CRIB-3mCitrine at cell tips is decreased, and CRIB-3mCitrine patches appear on cell sides (red arrowheads). However, in many cells (four right-hand columns), CRIB-3mCitrine appears to recover to cell tips within approximately 60 minutes (blue arrowheads). Scale bar, 5 µm. See also **Extended data Figure S6**.

**Figure 10.**
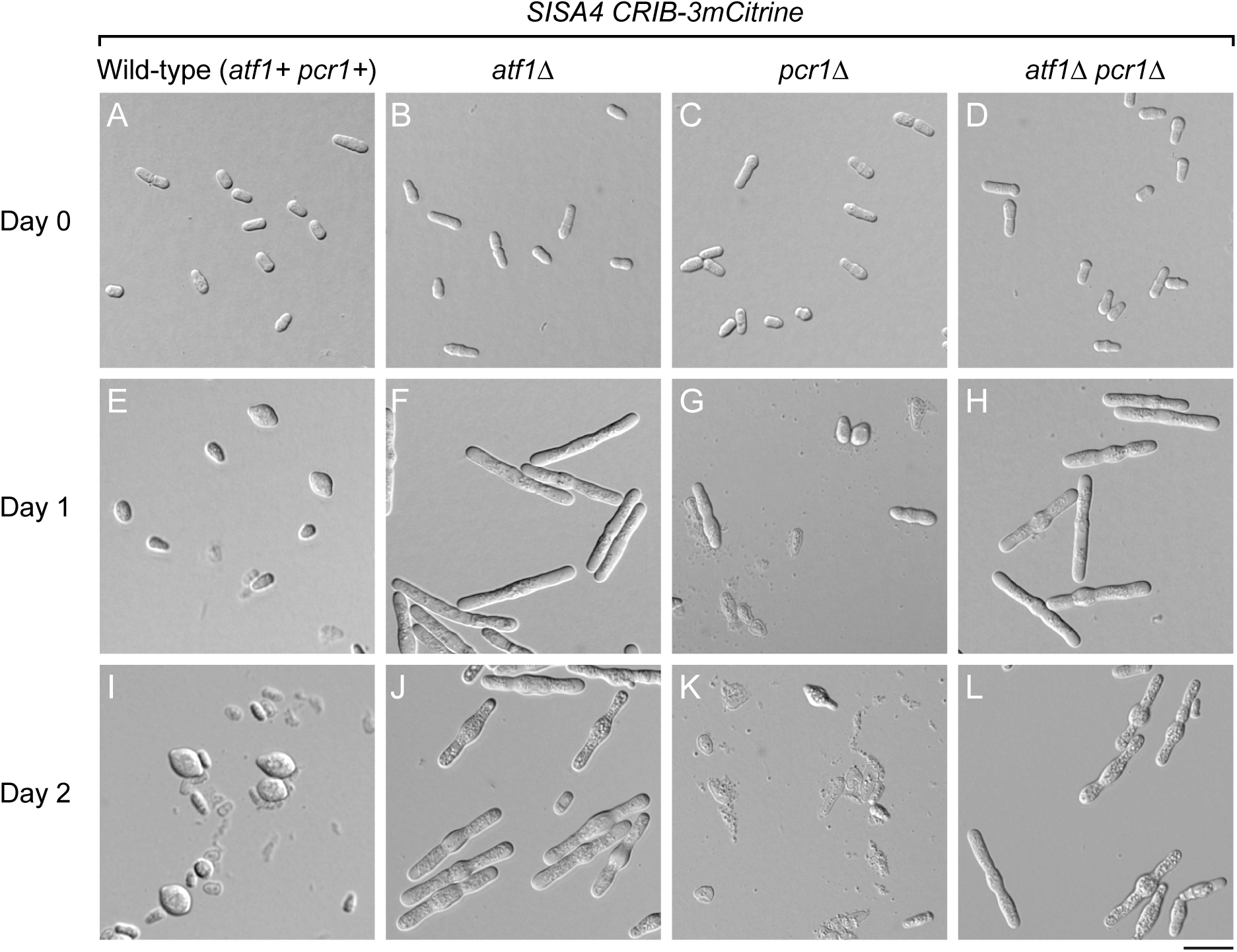
Deletion of Atf1 allows partial recovery of polarized growth in *SISA4* cells after Sty1 hyperactivation-induced polarity disruption. Cells from a *SISA4 CRIB-3mCitrine* background, containing the additional indicated mutations, at zero, one, and two days after plating to YE5S agar medium lacking AS-kinase inhibitor. Wild-type (i.e. *atf1+ pcr1+*) cells (A, E, I) are often swollen after one day, and more strongly swollen or lysed after two days. By contrast, *atf1Δ* cells (B, F, J) are significantly longer after one day, indicating resumption of polarized growth after initial polarity disruption (see **Figure 9**); they are also somewhat wider but do not divide. After two days, they do not lengthen further but continue to swell in central regions. They are less prone to lysis than wild-type cells but after two days often have a granular appearance. *pcr1Δ* cells (C, G, K) show a range of morphologies after one day (e.g. swelling, some lengthening, lysis) and are frequently lysed by two days. *atf1Δ pcr1Δ* double-mutant cells (D, H, L) resemble *atf1Δ* single mutants but may elongate less and swell more in central regions. After two days, they also have a granular appearance. Scale bar, 20 µm.

Atf1 normally functions by heterodimerizing with a second bZIP transcription factor, Pcr1 **(Kanoh *et al*., 1996; Kon *et al*., 1997; Wahls & Smith, 1994; Watanabe & Yamamoto, 1996)**. We therefore also imaged CRIB-3mCitrine localization after 1-NM-PP1 washout in *SISA4 pcr1Δ* cells (**Extended data Figure S7**). After washout, *SISA4 pcr1Δ* cells showed changes in CRIB-3mCitrine localization similar to *SISA4* (*pcr1+*) cells; unlike *SISA4 atf1Δ* cells, we did not observe any clear recovery of CRIB-3mCitrine to cell tips. Accordingly, *SISA4 pcr1Δ* cells were not highly elongated after one or two days of Sty1 hyperactivation; instead, they displayed a range of morphologies and were highly prone to lysis (**Figure 10C, G, K**).

The mechanism underlying the *SISA4 atf1Δ* phenotype after Sty1 hyperactivation remains unclear (see Discussion). Given that 1) *SISA4 atf1Δ* and *SISA4 pcr1Δ* showed different phenotypes after Sty1 hyperactivation, and 2) Atf1 and Pcr1 normally heterodimerize, we considered the possibility that the *SISA4 atf1Δ* phenotype could be due to inappropriate/abnormal function of Pcr1 when *atf1* is deleted. If this were the case, then one would expect that deletion of *pcr1* in *SISA4 atf1Δ* cells might revert the elongated morphology seen after long-term Sty1 hyperactivation. However, when we examined *SISA4 atf1Δ pcr1Δ* cells after one and two days of Sty1 hyperactivation, we found that they closely resembled *SISA4 atf1Δ* cells (**Figure 10F, H, J, L)**. This indicates that the *SISA4 atf1Δ* cell-polarity phenotype is independent of Pcr1 (see Discussion).

### Unusually low frequency of SISA4 progeny from crosses of SISA4 with wild-type (non-SISA) strains

For *SISA4* strains to be useful for studying processes regulated by Sty1, genetic crosses of *SISA4* with mutants or tagged strains of interest should be relatively easy to perform. Surprisingly, in crosses of *SISA4* with wild-type cells, we obtained an unexpectedly low frequency of *SISA4* progeny (**Figure 11**). Based on Mendelian segregation and *pyp1*-*pyp2* linkage, a cross between *SISA4* and wild-type (i.e. non-*SISA*) would be expected to produce about 12.5% total *SISA* progeny, with a *SISA4*:*SISA3* ratio of about 4:1 (i.e. 10% *SISA4* and 2.5% *SISA3*). However, in test crosses we routinely obtained only 0.3-0.8% *SISA4* progeny. Curiously, in these crosses *SISA3* progeny were produced at close to the expected frequency, such that *SISA3* progeny outnumbered *SISA4* progeny. This suggested that in a cross of *SISA4* with wild-type, there may be a specific problem in viability and/or germination of *SISA4* spores rather than in meiosis or spore formation more generally.

**Figure 11.**
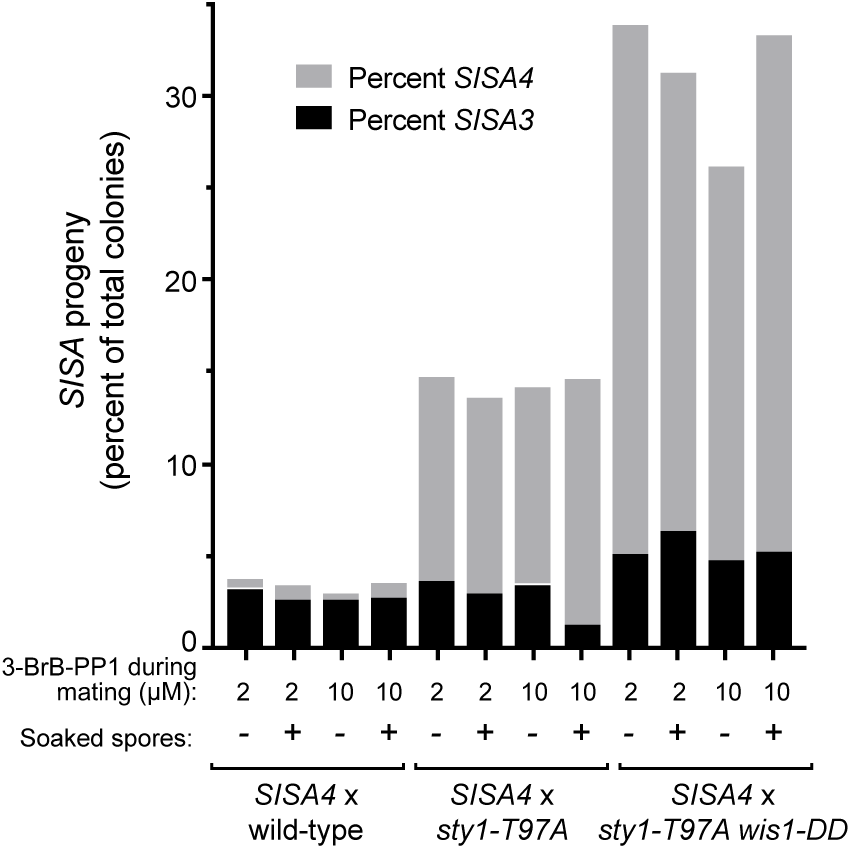
Frequency of *SISA4* and *SISA3* progeny from crosses of *SISA4* with non-*SISA* strains under conventional germination conditions. Spores from the indicated crosses were germinated on YE5S plates containing 2 µM 3-BrB-PP1 for 5 days at 32°C, before assaying *SISA* phenotypes. “3-BrB during mating” indicates concentration of 3-BrB-PP1 used on mating plates. “Soaked spores” indicates whether or not spores were soaked briefly in 50 µM 3-BrB-PP1 prior to germination. Expected Mendelian frequencies for *SISA4* and *SISA3* progeny were 10% and 2.5% (respectively) for *SISA4* x wild-type, 20% and 5% for *SISA4* x *sty1-T97A*, and 40% and 10% for *SISA4* x *sty1-T97A wis1-DD*. *SISA4* x wild-type crosses yielded far fewer *SISA4* progeny than expected. Total numbers of colonies scored for each data column were (from left to right): 434, 379, 391, 262, 774, 628, 624, 391, 475, 430, 415, 267.

Interestingly, in test crosses of *SISA4* with *sty1-T97A* cells, this problem mostly disappeared (**Figure 11**). Although the frequency of total *SISA* progeny (∼14%) remained lower than the expected Mendelian value (25%), the frequency of *SISA4* progeny increased considerably, and the *SISA4*:*SISA3* ratio was closer to the expected 4:1 value. Similarly, in test crosses of *SISA4* with *sty1-T97A wis1-DD*, the frequency of total *SISA* progeny (∼31%) also increased, although not to the expected Mendelian value (50%), and *SISA4*:*SISA3* ratios were also close to 4:1.

Based on the marked increase in *SISA4* progeny obtained from crosses of *SISA4* with *sty1-T97A* compared to crosses of *SISA4* with wild-type, we sought to rule out the possibility that our original *sty1-T97A* strain contained an additional (hypothetical) mutation, unlinked to *sty1*, that could be required for, or enhance, *SISA4* phenotype and/or viability. This scenario was formally possible because our original *sty1-T97A* strain was a precursor to both immediate parents of *SISA4* (**Figure 1B**), leading to a 12.5% chance that such a hypothetical mutation could also be present in the *SISA4* strain; if this were the case, then the hypothetical mutation would be guaranteed to be present in all progeny from a cross of *SISA4* to *sty1-T97A*, but in only half of the progeny from a cross of *SISA4* to wild-type.

To test for the existence of such a hypothetical mutation, we performed an outcrossing experiment (**Figure 12A**). We crossed our original *sty1-T97A* strain with wild-type cells and recovered 26 independent *sty1-T97A* strains. We then crossed each of the 26 *sty1-T97A* strains to *SISA4* and quantified the resulting *SISA4* progeny from the 26 crosses. If the original *sty1-T97A* strain were to contain a hypothetical mutation of the type proposed, then the frequencies of *SISA4* progeny from the 26 crosses would be expected to follow a bimodal distribution, in which about half of the crosses yield *SISA4* progeny near the expected Mendelian frequency (20%), while the other half of the crosses yield *SISA4* progeny at a much lower frequency, or not at all. By contrast, if there were no such mutation in the first place, then all 26 crosses would be expected to yield *SISA4* progeny near the expected Mendelian frequency (20%). The frequencies of *SISA4* progeny from the 26 crosses showed a unimodal distribution, with a mean value of 17.5% (**Figure 12B**). We further validated these results by repeating crosses for the three *sty1-T97A* strains that had yielded the lowest frequencies of *SISA4* progeny when crossed to *SISA4* (13.2%, 13.4%, and 14.0%); the repeat crosses yielded 16%, 20%, and 17% *SISA4* progeny (respectively), suggesting that these initial low frequencies were simply due to random variation in the first test.

**Figure 12.**
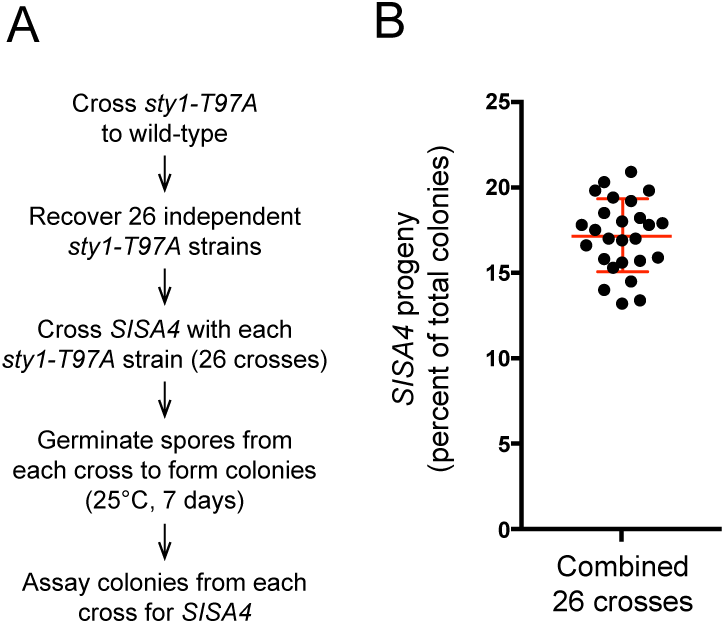
The four *SISA4* mutations are sufficient for the *SISA* phenotype. A. Outline of crosses and assay. See main text for details. **B.** *SISA4* progeny (as percent of total colonies) from crosses of *SISA4* with 26 independent *sty1-T97A* strains obtained after outcrossing. Line and error bars show mean and SD. Total number of colonies assayed in each of the 26 crosses ranged from 349 to 637.

We conclude that the original *sty1-T97A* strain used for *SISA4* construction does not contain any additional mutations affecting the *SISA4* phenotype, and that the four mutations *sty1-T97A*, *wis1-DD*, *pyp1Δ*, and *pyp2Δ* are sufficient for the *SISA4* phenotype. Our results further suggest that the low frequency of *SISA4* progeny in crosses of *SISA4* cells with wild-type cells may be due to germination issues associated with *SISA4* spores *specifically* derived from crosses of *SISA4* with wild-type (*sty1+*) cells. We speculate that this might occur via a mechanism involving cytoplasmic inheritance of wild-type Sty1 protein or mRNA during *SISA4* spore formation (see Discussion).

### Optimizing crosses of SISA4 with wild-type (non-SISA) strains

We next investigated whether we could improve the low frequency of *SISA4* progeny in crosses between *SISA4* and wild-type cells. The test crosses described above involved germinating spores at 32°C for 5 days on plates containing 2 µM AS-kinase inhibitor (in this case, 3-BrB-PP1) before assaying *SISA* phenotypes. We therefore tried germinating spores at 25°C for 7 days before assaying *SISA* phenotypes, and we also varied inhibitor concentration on germination plates. Under these conditions, the frequency of *SISA4* progeny from a cross of *SISA4* with wild-type increased significantly, especially at higher inhibitor concentrations, reaching up to 5-7.3% of total progeny (relative to a theoretical value of 10%; **Table 1**). In these experiments we also noticed that spores from a cross of *SISA4* with wild-type often germinated at different rates (**Extended data Figure S8**). Strikingly, the majority of *SISA4* progeny were found among the slowest-germinating colonies, including colonies that became visible by eye only after 7 days of incubation (**Table 1)**. Interestingly, this phenomenon appeared to be specific for crossing *SISA4* with wild-type; in a cross of *SISA4* with *SISA4* (in which all progeny are *SISA4*), most colonies germinated at normal rates **(Table 2**; **Extended data Figure S8).** We conclude that when *SISA4* is crossed with wild-type, *SISA4* progeny may have greater germination success at lower vs. higher temperatures, and, even then, they may take an unexpectedly long time to germinate (see Discussion). Accordingly, by controlling germination conditions and allowing sufficient time for colonies of the desired genotype to form, *SISA4* progeny can be recovered from crosses of *SISA4* with wild-type cells at suitable frequencies; the benefits of low temperature may be specific to germination, as we have found that vegetative growth of *SISA4* cells is better at 32°C than at lower temperatures. Based on these results, in the Methods section we present a detailed protocol for efficient recovery of *SISA4* progeny from crosses of *SISA4* cells with wild-type cells.

**Table 1.**
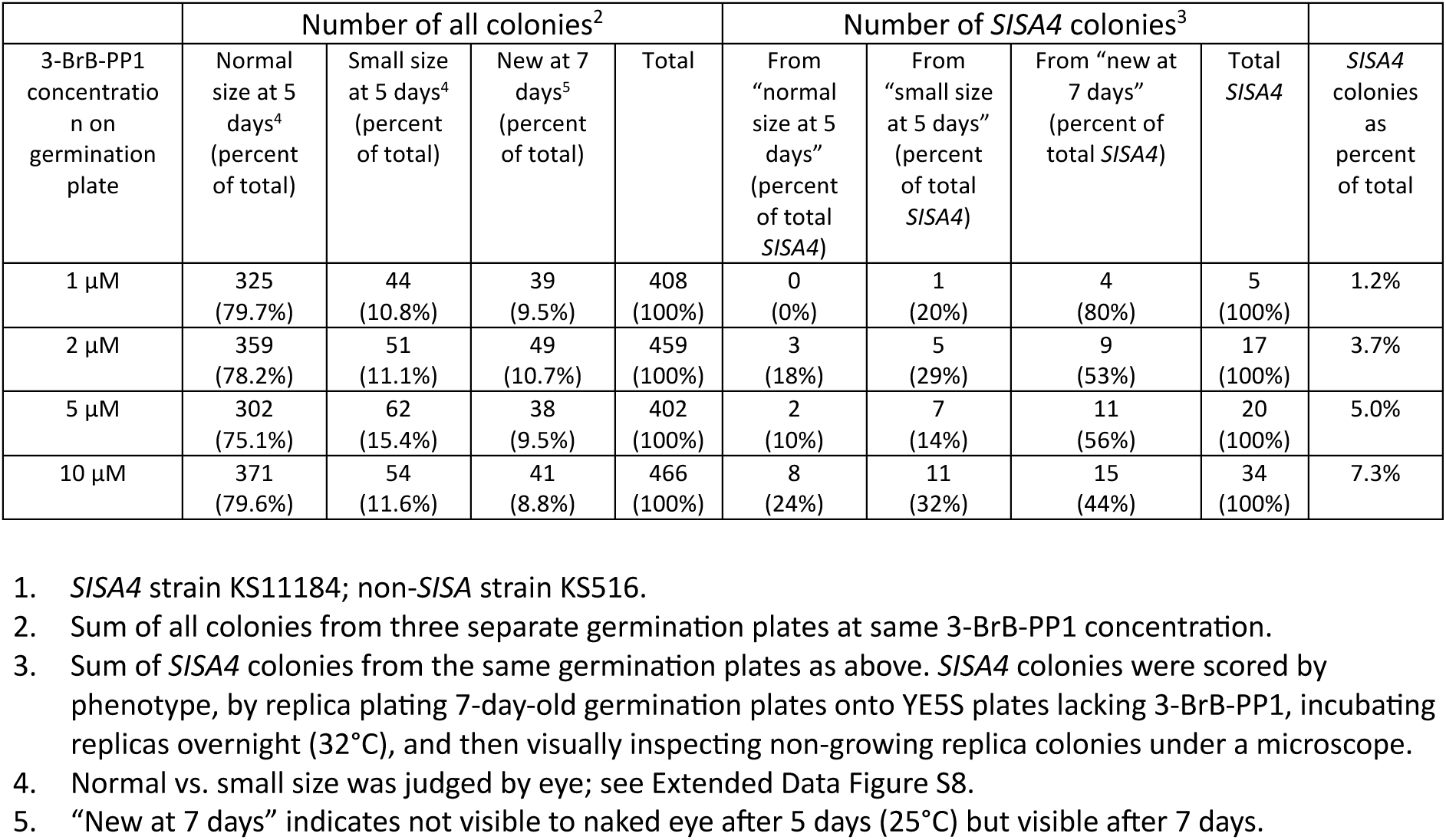
Size distribution of all colonies and SISA4 colonies in progeny from cross of SISA4 strain with non-SISA strain^1^ 1. *SISA4* strain KS11184; non-*SISA* strain KS516. 2. Sum of all colonies from three separate germination plates at same 3-BrB-PP1 concentration. 3. Sum of *SISA4* colonies from the same germination plates as above. *SISA4* colonies were scored by phenotype, by replica plating 7-day-old germination plates onto YE5S plates lacking 3-BrB-PP1, incubating replicas overnight (32°C), and then visually inspecting non-growing replica colonies under a microscope. 4. Normal vs. small size was judged by eye; see Extended Data Figure S8. 5. “New at 7 days” indicates not visible to naked eye after 5 days (25°C) but visible after 7 days.

**Table 2.** Size distribution of progeny from cross of *SISA4* strain with *SISA4* strain^1^ 1. *SISA4* strains KS11184 and KS11183. 2. Sum of colonies from three agar plates at same 3-BrB-PP1 concentration. All progeny are *SISA4*. 3. Normal vs. small size was judged by eye; see Extended data Figure S8. 4. “New at 7 days” indicates not visible to naked eye after 5 days (25°C) but visible after 7 days.

| 3-BrB-PP1 concentration on germination plate | Number of all colonies<sup>2</sup> |  |  |  |
| --- | --- | --- | --- | --- |
|  | Normal size at 5 days (percent of total) | Small size at 5 days<sup>3</sup> (percent of total) | New at 7 days<sup>4</sup> (percent of total) | Total |
| 1 μM | 316 (77.6%) | 51 (12.5%) | 40 (9.8%) | 407 (100%) |
| 2 μM | 447 (92.2%) | 27 (5.6%) | 11 (2.3%) | 485 (100%) |
| 5 μM | 384 (95.5%) | 9 (2.2%) | 2 (0.5%) | 395 (100%) |
| 10 μM | 426 (97.9%) | 7 (1.6%) | 2 (0.5%) | 435 (100%) |

## DISCUSSION

### SISA4 compared to original SISA strain

*SISA4* has several advantages compared to our original *SISA* strain, which was used in a limited set of experiments **(Mutavchiev *et al*., 2016)**. First, none of the four mutations in *SISA4* (*sty1-T97A*, *wis1-DD*, *pyp1Δ*, *pyp2Δ*) is linked to a selectable marker, making it easy to combine *SISA4* mutations with other mutations and/or tagged genes. By contrast, in our original *SISA* strain, both the CRIB-3mCitrine reporter and *pyp2* disruption were marked with *LEU2*, and both *pyp1* disruption and *wis1-DD* were marked with *ura4+* **(Mutavchiev *et al*., 2016)**. Second, in spite of the absence of selectable markers for *SISA4*, it is straightforward to identify and confirm *SISA4* strains both by phenotype (e.g. distinguishing *SISA4* from *SISA3* strains), and by genotype, through yeast colony PCR, including allele-specific PCR (see Methods). Third, phenotype reversion frequency of *SISA4* is several hundred-fold lower than that of the original *SISA* strain, due to modifications at the *wis1* locus. This improved genetic stability may be particularly useful in contexts such as genetic modifier screens. Furthermore, in this work we have determined optimal concentrations of AS-kinase inhibitors for use with *SISA4*, addressed issues of off-target effects, and established robust methods for crossing *SISA4* with wild-type strains. This provides a solid basis for future work with *SISA4* strains in microscopy, biochemistry, and proteomics experiments.

### Concentrations of AS-kinase inhibitors to use in experiments

In multiple assays, 3-BrB-PP1 and 1-NM-PP1 were equally potent for inhibition of Sty1-T97A *in vivo*, suggesting that these AS-kinase inhibitors can be used interchangeably. A 2 µM concentration of either inhibitor is sufficient for inhibition. Conversely, washout from 2 µM inhibitor to no inhibitor appears to be optimal for Sty1 hyperactivation in *SISA4* cells. Higher AS-kinase inhibitor concentrations might be helpful in specific applications, but for general use, the extra associated cost would not seem justified. Furthermore, higher inhibitor concentrations might lead to increased likelihood of off-target effects (see below). Our results also suggest that at low inhibitor concentrations (e.g. in the range of 0.2-0.5 µM) it may be possible to achieve mild or intermediate levels of Sty1 hyperactivation in *SISA4* cells, but we have not investigated this further. Based on cost, availability, and its overall wider adoption, we recommend using 1-NM-PP1 over 3-BrB-PP1 for general use with *SISA4*.

### AS-kinase inhibitor off-target effects

The evidence that inhibitors 3-BrB-PP1 and 1-NM-PP1 have modest off-target effects in fission yeast comes from the increased Lsd90-3mNG fluorescence seen in wild-type cells after AS-kinase inhibitor washout (**Figure 5**; **Figure 6**). We envision that during growth in the presence of AS-kinase inhibitors, inhibitor binding to off-target proteins leads to cells sensing and adapting to the presence of the inhibitors, and when inhibitor concentrations are later suddenly decreased (by washout), cells sense the decrease and activate a mild stress response, leading to increased *lsd90* expression via Sty1 and the CESR. While this is a speculative scenario, it is consistent with all our observations. Our results further suggest that cells can sense even micromolar-scale differences in inhibitor concentrations. Although the off-target interactors of AS-kinase inhibitors remain unknown, our inhibitor-swap experiments suggest that 3-BrB-PP1 and 1-NM-PP1 share the same key off-target interactors.

How should off-target effects be considered? In general, off-target effects of AS-kinase inhibitors *in vivo* have been thought to be minimal, provided that inhibitors are used at suitably low concentrations **(Zhang *et al*., 2013)**. For example, although a small number of wild-type mammalian protein kinases are known to be sensitive *in vitro* to certain AS-kinase inhibitors, including 1-NM-PP1 **(Bain *et al*, 2007; Bishop *et al*., 2000; Zhang *et al*., 2013)**, experimental evidence suggests that such kinases are unlikely to be significantly affected by micromolar or sub-micromolar AS-kinase inhibitor concentrations *in vivo,* where cellular ATP concentrations are typically 3-10 mM **(Zhang *et al*., 2013)**. Similarly, chemical genetic profiling of cell fitness in budding yeast has suggested that 500 nM 1-NM-PP1 has minimal off-target effects *in vivo* **(Hoon *et al*, 2008)**. At the same time, however, in previous work we found that adding a high (30 µM) concentration of 3-BrB-PP1 to wild-type cells led to a four-fold increase in expression of acid phosphatase Pho1 **(Tay *et al*, 2019)**. High concentrations of 1-NM-PP1 (50 µM) have also been suggested to have off-target effects in fission yeast *in vivo* **(Bohnert *et al*, 2020)**. Clearly, whether or not off-target effects are identified for a given AS-kinase inhibitor will depend both on the concentration of inhibitor used in experiments and on the type and sensitivity of the assays.

In this context, we note that the Lsd90-3mNG fluorescence assay is extremely sensitive. In *SISA4* cells, Lsd90-3mNG fluorescence increases several hundred-fold after washout of either 3-BrB-PP1 and 1-NM-PP1; by contrast, in wild-type cells, the corresponding increase is only 15-30-fold. Based on this, we think that although off-target effects of 3-BrB-PP1 and 1-NM-PP1 may exist in fission yeast, they can be considered to be minor relative to on-target effects, and they can be managed by using appropriate inhibitor concentrations and appropriate controls. Accordingly, while our imaging experiments showed that 1-NM-PP1 washout in *SISA4* cells leads to decreased CRIB-3mCitrine at cell tips, appearance of CRIB-3mCitrine patches on cell sides, and arrest of both cell elongation and cell-cycle progression, we found that 1-NM-PP1 washout in wild-type cells has absolutely no effect on CRIB-3mCitrine localization, cell elongation, or cell-cycle progression.

We also note that because washout from 5 µM AS-kinase inhibitor to no inhibitor leads to much higher Lsd90-3mNG fluorescence in *SISA4* cells than in wild-type cells, it might seem paradoxical that washout from 5 µM to 1-2 µM inhibitor leads to a greater increase in Lsd90-3mNG fluorescence in wild-type cells than in *SISA4* cells **(Figure 5)**. However, this can be explained by the dependency of increased *lsd90* expression on Sty1 kinase activity **(Chen *et al*., 2003; Rubio *et al*., 2021) (Extended data Figure S3C)**. That is, even if both wild-type and *SISA4* cells could sense the difference between, for example, 5 µM and 1-2 µM AS-kinase inhibitor concentrations, in *SISA4* cells Sty1 kinase activity would be inhibited by 1-2 µM AS-kinase inhibitor, thus preventing any consequent increase in *lsd90* expression.

### Disruption of the Cdc42 cell-polarity module after Sty1 hyperactivation in SISA4

The changes in localization of CRIB-3mCitrine seen in CHX-treated *SISA4* cells after 1-NM-PP1 washout indicate that new protein synthesis is not required for the disruption of cell polarity observed after Sty1 hyperactivation, consistent with the view that the changes driving polarity disruption after SAPK activation are mainly post-translational **(Mutavchiev *et al*., 2016)**. This in turn might seem to imply that proteins involved in stress-induced gene expression “downstream” of Sty1 should also not be involved in SAPK-driven disruption of cell polarity. However, our results with *SISA4 atf1Δ* cells suggest a somewhat more complex picture. After 1-NM-PP1 washout, *SISA4 atf1Δ* cells initially lose CRIB-3mCitrine from cell tips, but they can later recover it to cell tips and eventually become highly elongated relative to *SISA4* (*atf1+*) cells. This suggests that although Atf1 is not required for initial polarity disruption after Sty1 hyperactivation, it has a role in sustaining polarity disruption over longer periods. What this role is, however, remains unclear. Given our results from CHX treatment, it is not immediately apparent how the *SISA4 atf1Δ* phenotype after 1-NM-PP1 washout could be directly related to Atf1’s function in stress-induced transcriptional activation and the CESR. As *SISA4 atf1Δ* cells do not progress to cell division after 1-NM-PP1 washout, it seems likely that Sty1 remains hyperactivated in these cells. However, after two days of Sty1 hyperactivation, *SISA4 atf1Δ* cells do not elongate further but instead swell in the middle, more like *SISA4* (*atf1+*) cells (**Figure 10E ,F, I, J**); thus, it is possible that the polarity “recovery” seen in *SISA4 atf1Δ* cells is itself only transient.

Further investigation in this area may shed additional light on the function(s) of Atf1, which extend beyond Atf1’s widely-recognized role as a stress-induced transcriptional activator. For example, in the absence of stress and/or Sty1 activity (e.g. as would be the case for *SISA4* cells grown in 1-NM-PP1), Atf1 appears to act as a transcriptional repressor for at least some target genes **(Degols & Russell, 1997; Sanso *et al*., 2008)**, and it is also required for heterochromatin formation at the *mat* locus **(Fraile *et al*, 2022; Jia *et al*, 2004; Kim *et al*, 2004; Nickels *et al*, 2022; Sun *et al*, 2024).** In this context, it is possible that Atf1’s role in sustaining cell-polarity disruption might in fact be executed before rather than after 1-NM-PP1 washout. In addition to its functions related to gene expression, Atf1 is also important for Sty1 accumulation in the nucleus upon osmotic stress **(Gaits *et al*., 1998)**, and for growth at cold temperature, via a mechanism unrelated to the SAPK pathway **(Soto *et al*., 2002)**. Finally, it is also of interest that *SISA4 atf1Δ* and *SISA4 pcr1Δ* cells have distinct cell-polarity phenotypes after 1-NM-PP1 washout, even though Atf1 usually functions in a heterodimer with Pcr1. This finding may be consistent with previous work showing that *atf1Δ* and *pcr1Δ* cells do not have identical stress sensitivities; while both *atf1Δ* and *pcr1Δ* cells are similarly sensitive to heat and oxidative stress, *pcr1Δ* cells are less sensitive than *atf1Δ* cells to osmotic stress **(Sanso *et al*., 2008)**. In line with this, although most CESR genes are commonly regulated by Atf1 and Pcr1, a small number of CESR genes do not require Pcr1 for stress-induced expression **(Sanso *et al*., 2008)**.

### Germination of SISA4 after crosses of SISA4 with wild-type cells

A surprising finding from our experiments with genetic crosses was that the rate and/or efficiency of germination of *SISA4* spores is impaired when spores are derived from a cross of *SISA4* with wild-type cells but not when spores are derived from a cross of *SISA4* with *sty1-T97A* cells. How might this occur? We speculate that in a cross between *SISA4* and wild-type, a spore that is genotypically *SISA4* may receive wild-type Sty1 protein, or *sty1* mRNA, from the *sty1^+^* parent, via cytoplasmic inheritance from the zygote. Within such a *SISA4* spore, wild-type Sty1 protein could then become hyperactivated by *wis1-DD*—that is, even in the presence of AS-kinase inhibitor—with consequent effects on germination. We note that for this model to make sense, it may be necessary that wild-type Pyp1 and/or Pyp2 proteins (or mRNA), which provide negative feedback on Sty1 activity, would not be inherited to *SISA4* spores in the same way as wild-type Sty1; alternatively, Pyp1 and/or Pyp2 might not be as stable as Sty1, or they might simply be insufficient to counteract the effects of hyperactivated wild-type Sty1. While this model is speculative, the general principle that spore viability can be impaired as a result of mechanisms involving cytoplasmic inheritance is well established; for example, this is the basis for meiotic drive by *wtf* elements **(Hu *et al*, 2017; Nuckolls *et al*, 2017).**

## METHODS

### Fission yeast methods

Standard methods and media were used for fission yeast culture and crosses **(Ekwall & Thon, 2017; Forsburg & Rhind, 2006; Petersen & Russell, 2016).** Unless indicated otherwise, cells were grown in YE5S medium, using Bacto yeast extract (Becton Dickinson). For minimal medium, PMG (Pombe Minimal Glutamate, also known as EMMG or Edinburgh Minimal Medium using Glutamate; **(Petersen & Russell, 2016))** was used. Supplements for auxotrophic mutants were used at 150-175 mg/L unless indicated otherwise. Solid media were prepared using 2% Bacto agar (Becton Dickinson). AS-kinase inhibitors 3-BrB-PP1 (product code TRC-A602985; CAS no. 956025-99-3) and 1-NM-PP1 (product code TRC-A603003; CAS no. **221244-14-0**) were obtained from LGC Standards. Concentrated AS-kinase inhibitor stock solutions (50 mM 3-BrB-PP1 in methanol; 50 mM 1-NM-PP1 in DMSO) were stored long-term at -80°C and for at least several weeks at -20°C. For growth of *SISA4* and *SISA3* strains, liquid medium with added AS-kinase inhibitor could be used for at least 2-3 weeks (stored at room temperature), and solid medium with added AS-kinase inhibitor could be used for at least 2 months (stored at 4-5°C, protected from bright light). Genetic crosses were performed using SPA5S agar (with 45 mg/L supplements) for mating, and either tetrad dissection or random spore analysis for isolation of progeny **(Ekwall & Thon, 2017)**.

Apart from SISA4 and SISA3 strain construction (see below), gene-tagging and gene-deletion were performed using PCR and homologous recombination **(Bahler *et al*, 1998)**. Tagging of Lsd90 with 3xmNeonGreen used template plasmid pKS1935 (pFA6a-3xmNeonGreen:hphMX6) and oligonucleotides OKS4544 and OKS4545. Deletion of *atf1* coding sequence used template plasmid pKS108 (pFA6a-kanMX6) and oligonucleotides OKS2558 and OKS2559. Deletion of *pcr1* coding sequence used template plasmid pKS1724 (pFA6a-bsdMX6) and oligonucleotides OKS4989 and OKS4990.

For simplicity, in this work, we frequently use the term “wild-type” to refer to control strains that have wild-type alleles relative to the mutants shown in the same figure/panel. However, “wild-type” strains described in this way may also have additional auxotrophies and/or tagged genes. Full genotypes of all yeast strains used are indicated in the strain list in **Extended data Table S2**.

### SISA4 and SISA3 strain construction

*SISA4* (*sty1-T97A wis1-DD pyp1Δ pyp2Δ*) and *SISA3* (*sty1-T97A wis1-DD pyp1Δ*) strains were constructed by first making a *sty1-T97A pyp1Δ pyp2Δ* strain by CRISPR/Cas9 **(Torres-Garcia *et al*., 2020)** and, in parallel, a markerless *sty1-T97A wis1-DD* strain; these two strains were then crossed (**Figure 1A)**.

To construct CRISPR plasmids to delete *pyp1+* and *pyp2+*, we used the SpEDIT system, which contain SpCas9 and sgRNA sequences on a single plasmid **(Torres-Garcia *et al*., 2020)**. Candidate sequences for sgRNAs and homology repair templates were chosen using CRISPR4P **(Rodriguez-Lopez *et al*, 2016)**. To construct the *pyp1Δ* CRISPR plasmid, we annealed sgRNA oligonucleotides OKS4824 and OKS4825 and introduced the resulting double-stranded DNA into pLSB-Nat **(Torres-Garcia *et al*., 2020)** via GoldenGate reaction using BsaI. To construct the *pyp2Δ* CRISPR plasmid, we used an identical approach, using oligonucleotides OKS4830 and OKS4831. The resulting Cas9-sgRNA plasmids were named pKS2327 for *pyp1Δ* CRISPR and pKS2328 for *pyp2Δ* CRISPR. Homology repair (HR) templates were generated by annealing oligonucleotides OKS4826 and OKS4827 (for *pyp1Δ)* and OKS4834 and OKS4835 (for *pyp2*Δ) via their complementary 3’ regions and then extending 3’ ends with Phusion polymerase, to make double-stranded DNA **(Torres-Garcia *et al*., 2020)**. HR templates were designed to replace the entire *pyp1+* or *pyp2+* open-reading frame with a single “A” base **(Rodriguez-Lopez *et al*., 2016).**

To make *sty1-T97A pyp1Δ pyp2Δ*, strain KS7818 (*sty1-T97A*) was co-transformed by electroporation with pKS2328 and *pyp2Δ* HR template **(Torres-Garcia *et al*., 2020)**. In this and subsequent steps, cells were grown in the presence of AS-kinase inhibitor 3-BrB-PP1, although later this was found not to be strictly necessary until construction of the final quadruple-mutant strain *sty1-T97A wis1-DD pyp1Δ pyp2Δ* (subsequently termed “*SISA4*”). The resulting *sty1-T97A pyp2Δ* strain (KS11179) was co-transformed with pKS2327 and *pyp1Δ* HR template **(Torres-Garcia *et al*., 2020).** The resulting *sty1-T97A pyp1Δ pyp2Δ* strain was named KS11181.

To generate a markerless, untagged *wis1-DD* strain, strain KS7902 (*h- wis1DD:12myc::ura4+ leu1-32 ura4-D18*; Kazuhiro Shiozaki lab strain KS2081; kindly provided by Kazuhiro Shiozaki, Nara Institute of Science and Technology) **(Shiozaki *et al*., 1998)** was grown in bulk on a YE5S plate, and cells were then resuspended in YE5S and plated to PMG plates containing 150 mg/L leucine and 37.5 mg/L uracil and 0.2% 5-fluoroorotic acid (5-FOA) to select for loop-out excision. 5-FOA-resistant colonies were selected and screened for presence of *wis1-DD* and absence of *wis1+* sequences. The resulting *wis1-DD* strain was named KS9459. This was crossed to *sty1-T97A* strain KS11169 to generate *sty1-T97A wis1-DD* strain KS11171.

To make *SISA4* (*sty1-T97A wis1-DD pyp1Δ pyp2Δ*) and *SISA3* (*sty1-T97A wis1-DD pyp1Δ*) strains, KS11181 was crossed with KS11171 and tetrads dissected on YE5S plates containing 5 µM 3-BrB-PP1. After germination and colony formation, YE5S/3-BrB-PP1 plates were replica-plated to YE5S plates and incubated overnight. Colony-growth phenotypes were assayed by eye, and cell-morphology phenotypes were assayed by bright-field microscopy. Genotypes were determined by yeast colony PCR. Recovered *SISA4* strains were KS11183 and KS11184. Recovered *SISA3* strains were KS11186 and KS11187.

### Yeast colony PCR to check SISA4 strains

The four mutations in *SISA4* strains were routinely confirmed by yeast colony PCR, using a slight modification of methods of Akada et al. **(Akada *et al*, 2000)**. To prepare samples for PCR, fresh (i.e. day-old) yeast patches or colonies on plates (∼2-3 mm diameter; corresponding to about 8 mg wet weight) were picked and resuspended in microfuge tubes containing 50 µL TE buffer plus 0.25% SDS and heated to 95°C for 5 min. Tubes were then spun in a room-temperature microfuge for 5 min at 13,000 rpm, and ∼40 µL of supernatant recovered, avoiding contamination from the pellet. 1 µL of supernatant was used as template DNA in 25 µL PCR reactions. PCR reactions also contained: 1X PCR Buffer IV (1X is 75 mM Tris-HCl, pH 8.8, 200 mM ammonium sulfate, 0.1% Tween 20); 1.5 mM magnesium chloride; 0.2 mM each dNTP; 0.5 µM forward primer; 0.5 µM reverse primer; 1% (v/v) Triton X-100; ∼5 U *Taq* polymerase; ∼0.25 U *Pwo* polymerase; plus sterile deionized water to 24 µL). To detect *wis1-DD*, forward and reverse primers were OKS3898 (allele-specific for Wis1-DD) and OKS3951, respectively. To detect *pyp1Δ,* forward and reverse primers were OKS4856 and OKS4853. To detect *pyp2Δ,* forward and reverse primers were OKS4860 and OKS4861. To detect *sty1-T97A*, forward and reverse primers were OKS4971 (allele-specific) and OKS4974. To detect *sty1+*, forward and reverse primers were OKS4987 (allele-specific) and OKS4974. Primer sequences for PCR and expected PCR product sizes are listed in Extended data Table S3. To amplify *wis1-DD*, *pyp1Δ* and *pyp2Δ* sequences, the PCR program was: 94°C 3 min; 30 cycles of 94°C 10 sec, 52°C 30 sec, and 68°C 2 min; 68°C 5 min; 20°C hold. To amplify *sty1-T97A* and *sty1+* sequences, the PCR program was: 95°C 2 min; 30 cycles of 95°C 10 sec, 50°C 15 sec, and 72°C 2 min; 72°C 5 min; 20°C hold. All oligonucleotides used are listed in **Extended data Table S3**.

***Crossing SISA4 strains with wild-type (non-SISA) strains*: PROTOCOL**

**Notes**:

- This protocol describes how to recover optimal numbers of SISA4 progeny from crosses of *SISA4* strains with wild-type (non-*SISA*) strains, using random spore analysis.
- Although the protocol specifies AS-kinase inhibitor 1-NM-PP1, 3-BrB-PP1 can be used instead throughout, at the same concentrations (note that 3-BrB-PP1 stocks are prepared in methanol, while 1-NM-PP1 stocks are prepared in DMSO; see above).
- All yeast manipulations should be done under sterile conditions.
- For particularly complicated strain constructions involving crossing a strain of interest to *SISA4* (e.g. if tetrad analysis is required), it may be advantageous to first cross the strain of interest to *sty1-T97A* and then cross the resulting *sty1-T97A* progeny of interest to *SISA4,* as this can improve the germination efficiency of *SISA4* cells after a cross (see Results section, “Unusually low frequency of SISA4 progeny from crosses of SISA4 with wild-type (non-SISA) strain”).

**Day 0**

1. Prepare SPA5S agar plates containing 2 µM 1-NM-PP1 (added from stock solution just before pouring). 1 µM 1-NM-PP1 can also be used.

o *Although sty1Δ cells have been described to be sterile or mate poorly, we usually have no difficulty mating SISA4 cells in the presence of these concentrations of 1-NM-PP1. If mating efficiency is very low, it might be possible to improve mating by using slightly lower concentrations of 1-NM-PP1*.

2. Using fresh cultures, make a small but dense patch of each parent strain on appropriate solid media and grow overnight at 25°C.

**Day 1**

3. Put 10 µL sterile water in clean/sterile microfuge tube. Use P200 pipet tip to take a small amount (est. less than 1 µL; less than 1 mg) of each parent strain. Mix well by pipetting while pressing pipet tip against bottom of tube.

o *SISA4 cells can clump in culture, especially at lower temperature; this may impair mating with other parent strain. Pipetting against bottom of tube mitigates clumping better than vortexing*.

o *Good mixing of parent strains is important because inhibition of Sty1-T97A kinase activity by 1-NM-PP1 (which is necessary to prevent SISA4 lethality) is also deleterious to mating efficiency*.

4. Immediately pipet entire 10 µL mix onto SPA5S/1-NM-PP1 mating plate. Keep the area as small as possible.

o *If doing multiple crosses, do steps 3-4 for the first cross, then repeat for the second cross, etc. This will minimize re-clumping*.

5. Incubate mating plate 3 days at 25°C or 2 days at 28°C (if incubating at 28°C, all subsequent steps can be done one day earlier than specified in protocol).

**Day 4**

6. Prepare YE5S agar plates containing 2 µM 1-NM-PP1 (can be done in advance). Ensure that they are dry enough to accommodate 100 µL of diluted spore mixture during plating.

7. Dilute glusulase (Perkin Elmer, cat. no. NEE154001EA) 1:50 in sterile water (final glusulase 2% v/v). For each cross, use 2 µL of glusulase and 98 µL water. Scale up as needed. Spin in 4°C microfuge for 15 min at 13,000 rpm.

8. Recover diluted glusulase supernatant and discard any insoluble (brown) material.

9. For each cross, put 300 µL sterile water in microfuge tube. Use P200 tip to pick a small amount of mating mix (est. less than 1 µL; less than 1 mg) and add to tube.

o Pick from center of mating mix. This tends to have the highest number of spores.

10. Add diluted glusulase supernatant to mating mix (100 µL + 300 µL; final glusulase 0.5% v/v). Incubate for 2hr at 25°C.

o *Before adding glusulase to mating-mix suspension, you can look under microscope to judge the proportion of spores compared to vegetative cells. This is best done using a hemocytometer or multiwell slide. Using a conventional slide and coverslip may flatten cells/spores*.

o *If you want to save any undigested mating-mix suspension for reference, set aside a small volume of suspension before adding glusulase*.

o *If desired, the suspension can be vortexed occasionally during incubation (e.g. every 30 min)*.

11. Spin in room-temperature microfuge for 1 min at 13,000 rpm to pellet spores. Remove as much supernatant as possible. Resuspend in 1 mL sterile water.

o *Spore pellet is often not very tight. If you have many crosses, don’t spin too many tubes at once--if you do, the pellet may have loosened up in later tubes by the time you get to them, which makes it difficult to remove supernatant completely*.

o *You can also do a “double spin” to obtain a tighter pellet: spin in microfuge 1 min, then rotate tubes in rotor by 180° and spin again*.

o *Use a pipet to remove supernatant; don’t aspirate. We normally remove about half of the volume first, and then remove the rest carefully all the way down to the pellet. The goal is to remove as much glusulase as possible*.

12. Repeat spin, remove supernatant, and resuspend again in 1 mL sterile water.

13. Count in hemocytometer to determine the number of spores in 100 nL (nanoliters).

o *(optional) Some researchers include a wash with 30% ethanol to kill any remaining vegetative cells, although we do not find this to be necessary*.

14. Dilute spores in sterile water to obtain 400 spores per 100 µL. Plate 100 µL (i.e. 400 spores) on each YE5S/1-NM-PP1 plate. Spread with sterile glass beads.

o *Depending on efficiency of germination/viability, etc., total colonies from a cross between SISA4 and wild-type (sty1+ wis1+ pyp1+ pyp2+) strains may be only 35-50% of total spores plated, and total SISA4 colonies may be only 4-8% of colonies formed. Therefore, SISA4 colonies may be only 1-4% of total spores plated. Take this into account when deciding how many plates to use for each cross. Also take into account how many other markers are differentially present in each cross*.

o *Avoid plating too many spores per plate, because too many colonies can impair plate- and microscope-based phenotypic identification of SISA strains. Plating 400 spores per plate is based on assumption that only 35-50% of spores will form colonies (i.e. 140-200 colonies per plate). This is optimal for SISA4 phenotype identification*.

15. Incubate 7 days at 25°C.

o *Although vegetative growth of SISA4 strains is fine at 32°C, we find that germination of SISA4 spores is better at 25°C; this is especially important in crosses of SISA4 with wild-type (see main text)*.

o *We have found that in crosses of SISA4 with wild-type, SISA4 spores are over-represented among the smallest colonies (i.e. because they frequently germinate very slowly). Therefore, shorter incubation times can lead to a disproportionate loss of SISA4 progeny colonies*.

**Day 9 (optional)**

16. Examine germination plates and use a fine-tip marker pen to circle any colonies that are smaller than normal.

o *As mentioned above, SISA4 spores—especially in crosses of SISA4 with wild-type—frequently germinate very slowly compared to other genotypes. Identifying the smallest colonies in advance can help when screening for SISA4 phenotypes on Day 12*.

**Day 11**

17. (optional) Examine germination plates and use a fine-tip marker pen to circle any new (tiny) colonies that have appeared between Day 9 and Day 11.

o *As mentioned above, identifying smallest colonies in advance can help when screening for SISA4 phenotypes on Day 12*.

o *If you are curious, you can use a different color pen compared to that used on Day 9 to distinguish new colonies identified on Day 11 from small colonies identified on Day 9*.

18. (optional) Count total colonies on representative germination plates.

o *This is useful if you want to measure frequency of recovery of SISA4 progeny relative to expectation*.

o *Count by marking each colony with a fine-tip marker* pen *while simultaneously clicking handheld counter. Counting is sometimes easier using a light-box, if one is available*.

19. Replica-plate each germination plate (donor plate) to a well-dried YE5S plate and (optional) to a YE5S plate containing 2 µM 1-NM-PP1.

o *For replica plating we use Whatman No. 1 filter papers (150 mm circles; cat. no. 1001-150) and a homemade cylindrical replica-plating block. Cool, dry plates give best replica plating. Use two filter papers for each replica-plating*.

o *After pressing donor plate on filters, but before making replica(s), press the (clean) lid of the donor plate onto the filters to remove excess cells from the filters. This “lid clean-up” step also helps to spread out smaller colonies*.

o *(optional) Because you will need to recover viable SISA4 strains from a YE5S/1-NM-PP1 plate after screening, it can be helpful to have a YE5S/1-NM-PP1 recipient plate in addition to the original donor plate. You can also use this replica to compare with the replica made to YE5S*.

o *(optional) After making replicas, do a further clean-up of the replicas by pressing recipient plate gently onto a single fresh filter paper on the block. This can be helpful if there is uneven cell density on the replicas*.

20. Incubate replicas 1 day at 32°C.

o *The difference between growth and non-growth (or poor growth) is greater at 32°C than at 25°C*.

o *(optional) If you plan to recover SISA4 strains from the donor plate, it can be helpful to incubate the donor plates for one more day as well, to increase mass of donor colonies*.

**Day 12**

21. Identify non/poorly-growing colonies on YE5S replicas by eye, and circle with fine-tip marker pen.

o *Growth is assessed by density of cells in the replica. When there is no growth at all, it can be difficult to “see” a non-growing colony, especially if it is small. For optimal assessment, you may need to adjust lighting conditions and/or angle of viewing of the plate. We normally assess growth by looking at the agar, without the lid (i.e. not looking through plastic); in normal lab conditions, contamination is unlikely*.

o *If you labelled small colonies on germination plates, you can use these as reference to check corresponding replicas*.

22. Assay cell phenotypes of non/poorly-growing colonies by light microscopy (bright field illumination; 10X or 20X objective)

o *Nearly all non/poorly-growing colonies will be SISA4 or SISA3; these can be distinguished from each other by shape*.

o *We use an upright yeast-plate microscope (with long working-distance objectives) to image plates upside down, and during visual screening we directly label colonies on plates (with* fine-tip marker pen*) as “S4” (SISA4) or “S3” (SISA3)*.

o *If no suitable microscope is available, yeast colony PCR can be used to verify mutations, using appropriate primers (see yeast colony PCR methods, above)*.

23. Recover SISA strains of interest from YE5S/1-NM-PP1 donor plate or from YE5S/1-NM-PP1 replica.

24. Streak recovered strains onto YE5S/1-NM-PP1 plates to obtain single-colony clones. Incubate 2-3 days at 32°C to obtain colonies.

25. (optional) Return YE5S replicas to 32°C and incubate 1 day at 32°C.

**Day 13 (optional)**

26. If on Day 12 you returned YE5S replicas to 32°C for 1 additional day of incubation, re-examine non/poorly-growing colonies on the YE5S replicas from Day 12.

o *This can be useful if you found it difficult to distinguish SISA4 from SISA3 colonies on Day 12. After an additional day of incubation, SISA3 colonies tend to be visibly denser than SISA4 colonies*.

o *If you do this, continue by streaking recovered strains as in Step 24, and add one day to all subsequent steps*.

**Day 14/15**

27. Replica-plate YE5S/1-NM-PP1 streak plates to YE5S plate, and incubate replicas 1 day at 32°C

**Day 15/16**

28. Confirm SISA phenotype of single-clone colonies by microscopy.

29. If desired, perform yeast colony PCR to verify mutations, using appropriate primers.

30. Make patches of appropriate strains for subsequent storage/freezing.

### Phenotype reversion assay

To assay *SISA* phenotype reversion, we used original *SISA* (KS8226) and *SISA4* (KS11051) strains expressing the Cdc42-GTP reporter CRIB-3mCitrine **(Mutavchiev *et al*., 2016)**, because the original *SISA* strain (KS8226) was constructed with CRIB-3mCitrine in the strain background. Strains were streaked on YE5S plates containing 2 µM 1-NM-PP1 to allow single cells to form “precursor” colonies within which revertants could arise at random (i.e. under non-selective conditions). For each strain, ten precursor colonies were then harvested separately. Each precursor colony was resuspended in 200 µL YE5S, and the total number of cells in each suspension was determined by hemocytometer counting. For each suspension, half of the suspension (100 µL, ranging from 1.2x10^6^ to 4.2x10^6^ cells, depending on the precursor colony) was plated to a single YE5S plate and incubated for 3 days at 32°C. Only half of the harvested suspension was plated because some of the suspension was used for hemocytometer counting. The number of colonies on each plate was then counted; because *SISA*/*SISA4* cells normally cannot grow on YE5S plates lacking AS-kinase inhibitor, the number of colonies on each YE5S plate should be equal to half of the total number of revertant cells present within the harvested precursor colony.

A simple formula to estimate *p*, the probability of a revertant arising (per cell per generation) can be derived as follows: We make the simplifying assumptions that 1) *p* is very small, 2) reversion events are independent, and 3) revertant cells can divide and divide at the same rate as non-revertant cells. Let *g* be the number of generations to make a precursor colony from a single cell, such that the total number of cells *N* in the precursor colony is *2^g^.* Let *R* be the number of revertant cells present in the precursor colony. Now consider an individual generation *i* of precursor colony growth. In any given generation *i*, there will be *2^i^* cells. Therefore, the number of new revertants in generation *i* will be *p2^i^*. Next, consider that a new revertant cell appearing in generation *i* can divide and give rise to progeny. Therefore, after *g* generations, a revertant cell that first appeared in generation *i* will have given rise to *2^(g-i)^* revertant cells in the precursor colony. Accordingly, each generation *i* should give rise to a total of *p2^i^2^(g-i)^ = p2^g^* revertant cells in the precursor colony. Because the precursor colony had *g* generations of growth, the scenario described above for one individual generation *i* can occur *g* times. Therefore, the total number of revertant cells in the (final) precursor colony should be *R* = *gp2^g^.* From this, we obtain *p* = *R*/*g2^g^*= *R/gN.* Because *N* = 2*^g^*, we can write *g* = ln*N*/ln2. Therefore, *p* = *R*/((ln*N*/ln2)*N*) = Rln2/(*N*ln*N*). From this it can be seen that for two precursor colonies with equal numbers of cells (i.e. equal *N*) the fold-difference in *p* is directly proportional to the fold-difference in *R*, with no other factors contributing. From counting phenotype revertants **(Figure 2A),** we estimate *p* in the original *SISA* strain to be approximately once per 200,000 cells per generation. Because the number of revertants in *SISA4* was so low (often zero), it is more difficult to estimate *p* for *SISA4*; however, a generous estimate is in the range of once per 100,000,000 cells per generation, i.e. up to 500 times lower than for the original *SISA* strain.

### Time-lapse fluorescence microscopy

For live-cell time-lapse fluorescence microscopy of CRIB-3mCitrine, preparation of samples and imaging was performed essentially as described previously **(Mutavchiev *et al*., 2016)**; minor differences are described here. Cells were grown to log phase in YE5S liquid medium containing 2 µM 1-NM-PP1 at 25°C, mounted on 4-chamber microslides (Ibidi; 80427) that were pre-coated with 1 mg/mL soybean lectin (Sigma; L1395), and covered with approximately 0.5 mL growth medium. Imaging was performed on a Nikon TE2000 microscope (100x/1.45 NA Plan Apo objective) using a Yokogawa CSU-10 spinning disk confocal module with an iXon+ Du888 EMCCD camera (Andor), controlled by Metamorph software (Molecular Devices). A free, open-source alternative is µManager (Edelstein *et al*, 2014). Microscope temperature was maintained at 25°C in an environmental chamber. Imaging used a 50 mW 488 nm laser set at 20% laser power and a 488/594 dual-band dichroic filter. Timepoints were acquired at 5-min intervals. For each timepoint, 11 Z-sections were collected, with 0.6 µm spacing and 120 ms exposure per Z-section. Under these conditions there was negligible apparent photobleaching. For experiments involving 1-NM-PP1 washout, washout was performed by removing growth medium from the microslide chamber using a 1 mL transfer pipet with narrow tip (Fisher cat. no. 13469118) and replacing medium with 0.5 mL of medium lacking 1-NM-PP1. This was repeated 5 additional times to ensure complete washout of 1-NM-PP1. For mock wash experiments, growth medium was replaced with medium containing 1-NM-PP1. Because a complete set of 6 washes could take up to 10-12 minutes, the image acquisition program was paused during washout (and mock wash), leading to a small discontinuity in the time intervals around the time of washout. In the figures shown, “-5 min” represents the last timepoint prior to starting washout, and the “0 min” timepoint represents the first time point after completion of washout. For experiments involving cycloheximide (CHX), CHX was added to growth medium 30 minutes before washout at a final concentration of 100 μg/mL (from a 10 mg/mL stock in water); CHX was also present in growth medium after washout.

### Cell length at septation assay

To measure cell length at septation, cells were grown at 30°C for 48 hours in YE5S or PMG liquid medium (with appropriate supplements) containing different concentrations of 3-BrB-PP1 and 1-NM-PP1, maintaining log-phase growth throughout. Upon reaching cell density ∼1 × 10⁷ cells/mL, 900 µL of cell culture was fixed by addition of 100 µL formalin solution (37% Formaldehyde solution, Sigma, cat. no. F8775) and incubated on ice for 10 min. Cells were then washed twice with PBS and resuspended in 20 µL of PBS. Fixed cells were stained with Calcofluor (Fluorescent Brightener 28, Tinopal UNPA-GX, Sigma, cat. no. F3543). A 5 mg/mL stock solution of Calcofluor was prepared in DMSO and diluted 1:100 in PBS. To stain cells, this diluted stock was mixed with an equal volume of fixed cell suspension. Stained cells were imaged on multispot coated slides (Hendley-Essex, 12-well 9mm slide, cat. no. SM012) to prevent flattening of cells between slide and coverslip. Images were acquired using a Zeiss AxioImager Z1 microscope with a 100× oil immersion objective (alpha Plan-Apochromat 1.46 NA), using the DAPI channel. Images were processed and analyzed using Fiji/ImageJ, and the length of septating cells was measured manually by drawing a line along the long axis of the cells. Approximately 100 cells were measured per strain per condition. To ensure unbiased measurements, data were blinded during measurements.

### Flow cytometry

Flow cytometry experiments used cells containing an *lsd90-3xmNeonGreen* fluorescent reporter. For measurements of Lsd90-3xmNeonGreen expression after washout of AS-kinase inhibitors, cells were grown at 30°C for 48 hours in YE5S liquid medium containing either 2 µM or 5 µM 3-BrB-PP1 or 1-NM-PP1, with dilution to maintain log-phase growth throughout. Upon reaching cell density ∼1 × 10⁷ cells/mL, existing growth medium was removed from 1 mL of cells either by brief room temperature microcentrifugation (30 sec, 4k rpm) or by vacuum-filtration (Millipore, mixed cellulose ester, 25 mm, 0.025 µm pore size, cat. no. VSWP02500), and cells were then resuspended in fresh medium containing different concentrations of 3-BrB-PP1 or 1-NM-PP1 or in fresh medium alone. Cells were then grown for a further 3 hr after washout, to allow Lsd90-3xmNeonGreen expression, and then processed for flow cytometry (see below). Inhibitor swap experiments were performed similarly. For measurements of Lsd90-3xmNeonGreen expression at steady-state, cells were grown at 30°C for 48 hours in YE5S liquid medium containing either 2 µM or 5 µM 3-BrB-PP1 or 1-NM-PP1, or no inhibitor at all, maintaining log-phase growth throughout. Upon reaching cell density ∼1 × 10⁷ cells/mL, cells were directly processed for flow cytometry.

To process cells for flow cytometry, 100 µL of culture was diluted with 900 µL of the exact same type of medium in which cells were already growing (e.g. with or without AS-kinase inhibitors, at specific concentrations), to obtain an appropriate cell density for analysis. Flow cytometry was performed using an Attune NxT Flow Cytometer (ThermoFisher) to measure mNeonGreen fluorescence in the BL1 channel (510/10 bandpass filter). Photomultiplier tube (PMT) voltages were adjusted to maintain signals within the linear detection range, with the following settings: BL1 - 280 V, FSC - 200 V, and SSC - 340 V. Data were analyzed using FlowJo software (BD Biosciences), using gating strategies to exclude debris, doublets (based on forward and side scatter signals), and other contaminants, ensuring the analysis focused solely on single, viable cells. Approximately 100,000 events were analyzed per condition. Free, open-source alternatives to FlowJo include EasyFlow (https://antebilab.github.io/easyflow/) and EasyFlowQ (https://ym3141.github.io/EasyFlowQ/) **(Ma *et al*, 2024)**.

## Supporting information

Combined supplemental extended data

## DATA AVAILABILITY

### Underlying data

Edinburgh DataShare (University of Edinburgh Research Data Service): A chemical-genetic approach for stress-independent activation of the fission yeast stress-activated protein kinase pathway. https://doi.org/10.7488/ds/8147

This project contains the following underlying data:

- SISA4_SISA3_DIC.zip. ZIP file containing DIC microscopy data of *SISA4* and *SISA3* strains in the presence and absence of AS-kinase inhibitor 1-NM-PP1.
- Septation_length.zip. ZIP file containing fluorescence microscopy data of cell septation length.
- CRIB3mCitrine_video.zip. ZIP file containing fluorescence videomicroscopy data of Cdc42-GTP reporter CRIB-3mCitrine.
- SISA4_atf1D_pcr1D_DIC.zip. ZIP file containing DIC microscopy data of *SISA4* strains with additional mutations, in the absence of AS-kinase inhibitor.
- README.txt. Text file describing the structure of the repository.

All image data are in .tif format and can be opened using free open-source software FIJI **(Schindelin *et al*., 2012)**.

### Extended data

Edinburgh DataShare (University of Edinburgh Research Data Service): A chemical-genetic approach for stress-independent activation of the fission yeast stress-activated protein kinase pathway. https://doi.org/10.7488/ds/8147

- Sawin_et_al_combined_extended_data.pdf. PDF file containing all extended data figures and tables (supplementary data).
- README.txt. Text file describing the structure of the repository.

Data are available under the terms of the Creative Commons Zero “No rights reserved” data waiver (CC0 1.0 Public domain dedication).

## COMPETING INTERESTS

No competing interests were disclosed.

## GRANT INFORMATION

This work was supported by Wellcome [210659; Wellcome Trust Investigator Award in Science to KES]; [203149; core funding for the Wellcome Centre for Cell Biology]; [226791; the Wellcome Discovery Research Platform for Hidden Cell Biology]; [218470; Wellcome-funded 4 year PhD Programme in Integrative Cell Mechanisms (iCM) and an iCM Transition Award to DM], by the UK Biotechnology and Biological Sciences Research Council (BBSRC) [BB/M010996/1; BBSRC/EASTBIO PhD studentship to AK] and by the Darwin Trust of Edinburgh [PhD studentships to AG, BB, AIR-R and MLS].

## ACKNOWLEDGEMENTS

We thank members of our laboratory for discussions and Andreas Fellas and Alison Pidoux for advice on CRISPR-Cas9 methods for fission yeast. We gratefully acknowledge support from the Wellcome Discovery Research Platform for Hidden Cell Biology Light Microscopy Core and technologists David Kelly and Toni McHugh. For the purpose of open access, the authors have applied a Creative Commons Attribution (CC BY) licence to any Author Accepted Manuscript version arising from this submission.

## Notes

### Competing Interest Statement

The authors have declared no competing interest.

