## Supplementary material for "A chemical-genetic approach for stress-independent activation of the fission yeast stress-activated protein kinase pathway": Combined supplemental extended data

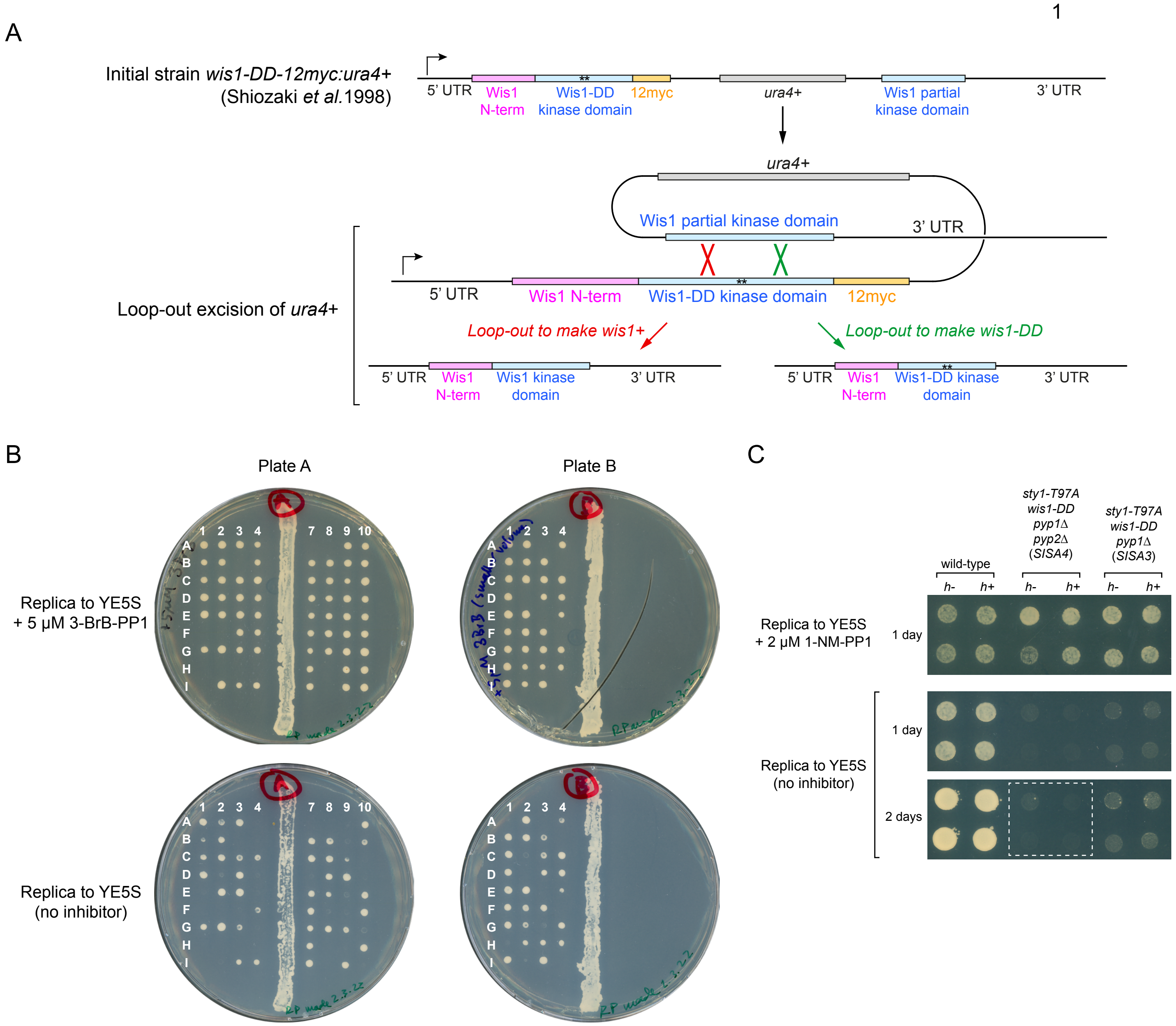


**Extended data Figure S1. Additional aspects of *SISA4* and *SISA3* strain construction.**

**A.** Generation of unmarked *wis1-DD* strain. The initial strain, made by Shiozaki et al. (1998), contained *wis1DD-12myc:ura4+* strain at the *wis1* locus and was constructed using "ends-in" recombination of a linearized *ura4+* plasmid containing a partial Wis1 kinase domain with activating mutations S469D and T473D (asterisks) and a C-terminal 12myc tag (see Shiozaki et al. 1998) for details). Spontaneous "loop-out" excision of *ura4+* by homologous recombination generates either *wis1+* or *wis1-DD* strains (both lacking 12myc tag), depending on whether recombination occurs 5' or 3' to the DD mutations (shown in diagram as red and green, respectively). Excision of *ura4+* allows cells to grow on minimal medium containing uracil plus 5-fluoroorotic acid (5-FOA).

**B.** Identification of *SISA* mutants by replica-plating from plates containing 5 μM AS-kinase inhibitor 3-BrB-PP1 to YE5S plates with and without inhibitor, followed by overnight incubation. All strains that failed to grow after replica-plating to YE5S without inhibitor were later identified to be either *sty1-T97A wis1-DD pyp1Δ pyp2Δ* (*SISA4*) or *sty1-T97A wis1-DD pyp1Δ* (*SISA3*). Colony labeling scheme corresponds to that shown in Extended data Table S1.

**C.** Replica plating of *SISA4* and *SISA3* strains to YE5S plates containing 2 μM AS-kinase inhibitor 1-NM-PP1 and to YE5S plates without inhibitor. For each strain, two duplicates are shown for *h-*, and two for *h+*. Although neither *SISA4* nor *SISA3* strains grow well after replica plating, *SISA4* colonies typically show slightly lower cell density than *SISA3* colonies, particularly 2 days after replica plating (dashed lines). This is usually easily identifiable by eye.

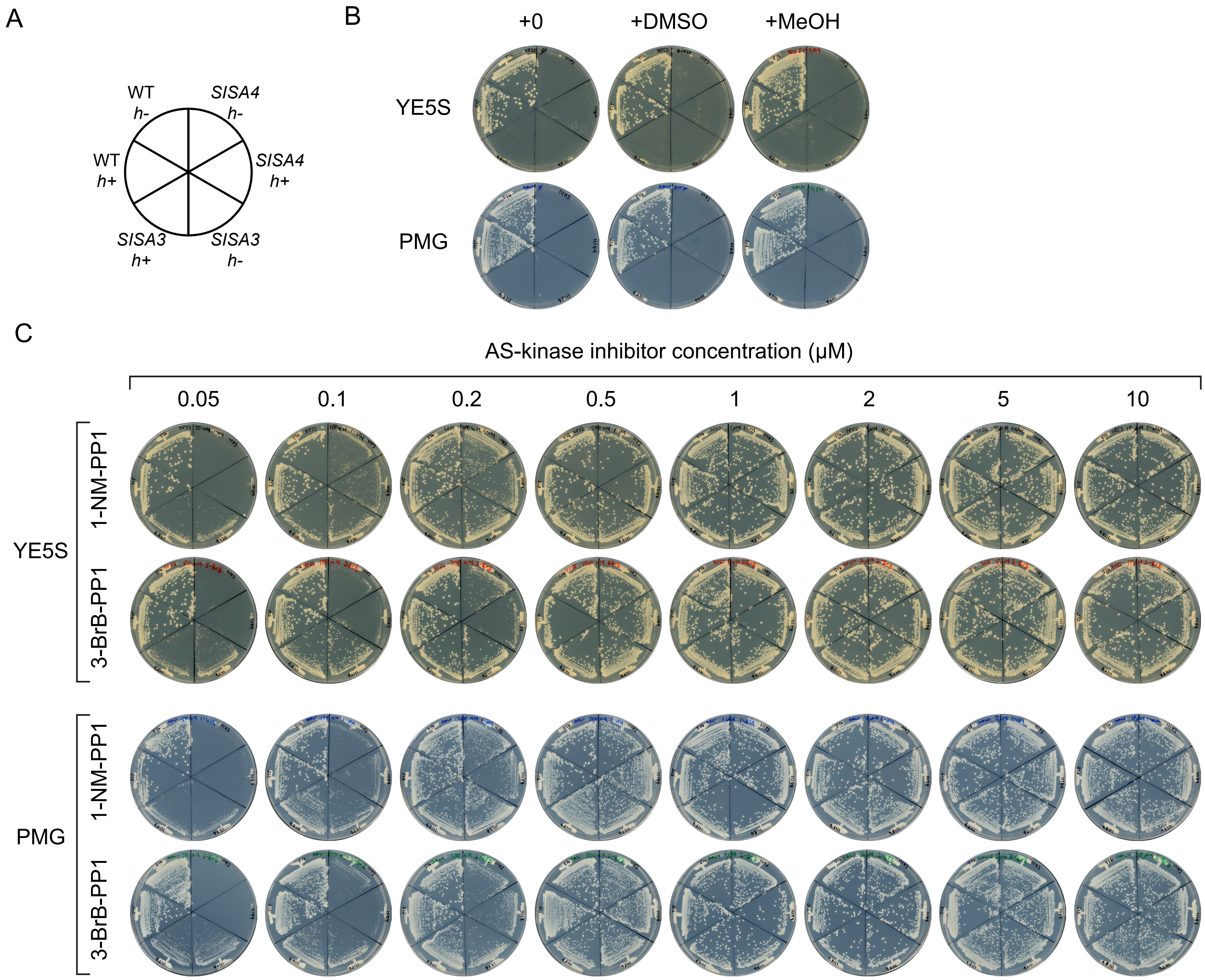


**Extended data Figure S2. AS-kinase inhibitors 1-NM-PP1 and 3-BrB-PP1 have similar effects on growth of *SlSA4* cells and similar effects on growth of *SlSA3* cells.**

**A.** Plan of the indicated strains streaked in B and C.

**B.** Colony formation on rich (YE5S) and minimal (PMG) agar plates with nothing added, or with solvents for AS-kinase inhibitors added. DMSO is used as solvent for 1-NM-PP1; methanol (MeOH) is used as solvent for 3-BrB-PP1. Plates were scanned after 3 days' growth at 30°C. Images of plates marked "+0" are copied from Figure 1.

**C.** Colony formation on rich and minimal agar plates at the indicated concentrations of AS-kinase inhibitors. Plates were scanned after 3 days' growth at 30°C. Measurements of colony diameters from these plates are shown in Figure 3.

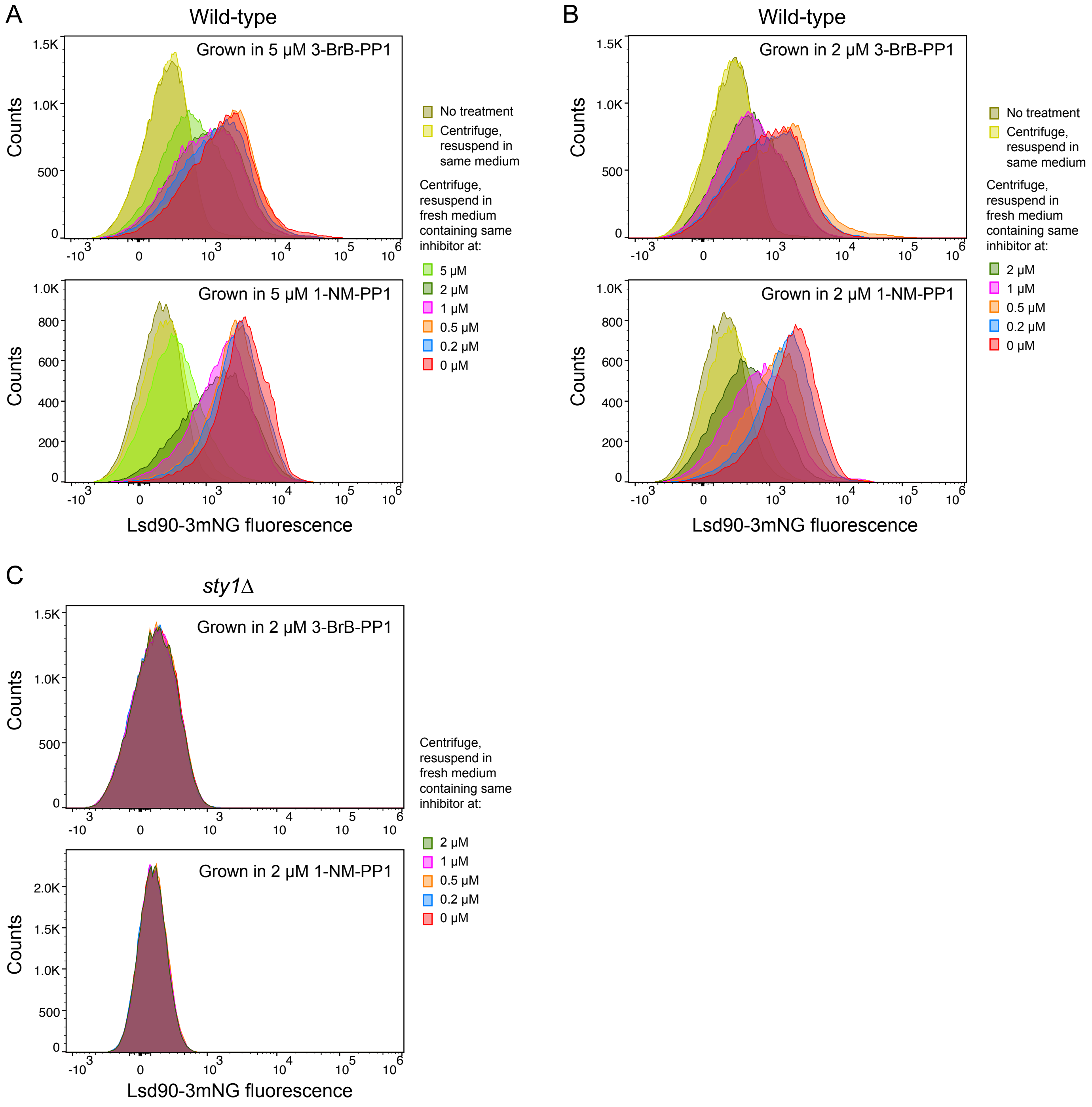


**Extended data Figure S3. Off-target effects leading to increased Lsd90 expression after AS-kinase inhibitor washout in wild-type cells depend on Sty1.**

**A, B.** Flow cytometry analysis of Lsd90-3xmNeonGreen (Lsd90-3mNG) expression in wild-type (non-*SISA*) cells three hours after centrifugation-based washout from growth medium containing (A) 5 μM or (B) 2 μM 3-BrB-PP1 or 1-NM-PP1 into medium containing different concentrations of the same inhibitors, or no inhibitor (0 μM).

**C.** Flow cytometry analysis of Lsd90-3mNG expression in *sty1*Δ cells after centrifugation-based washout from medium containing 2 μM 3-BrB-PP1 or 1-NM-PP1 into medium containing different concentrations of the same inhibitors, or no inhibitor (0 μM).

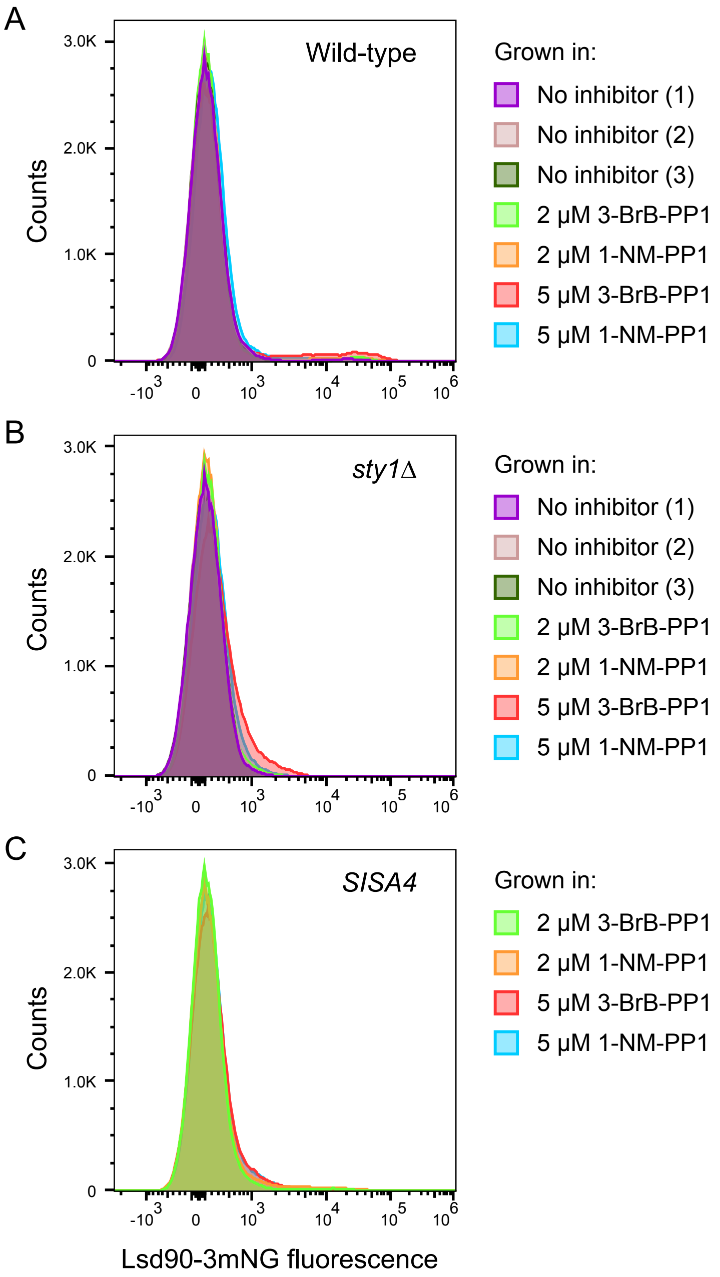


**Extended data Figure S4. AS-kinase inhibitors do not affect basal levels of Lsd90 expression.**
Flow cytometry analysis of Lsd90-3xmNeonGreen (Lsd90-3mNG) expression in (A) wild-type cells, (B) *sty1*Δ cells, and (C) *SISA4* cells grown in different steady-state concentrations of 3-BrB-PP1 and 1-NM-PP1, or in the absence of inhibitor (note that *SISA4* cells cannot be grown in the absence of inhibitor). For "no inhibitor" conditions, three independent cultures were analyzed for both wild-type and *sty1*Δ.

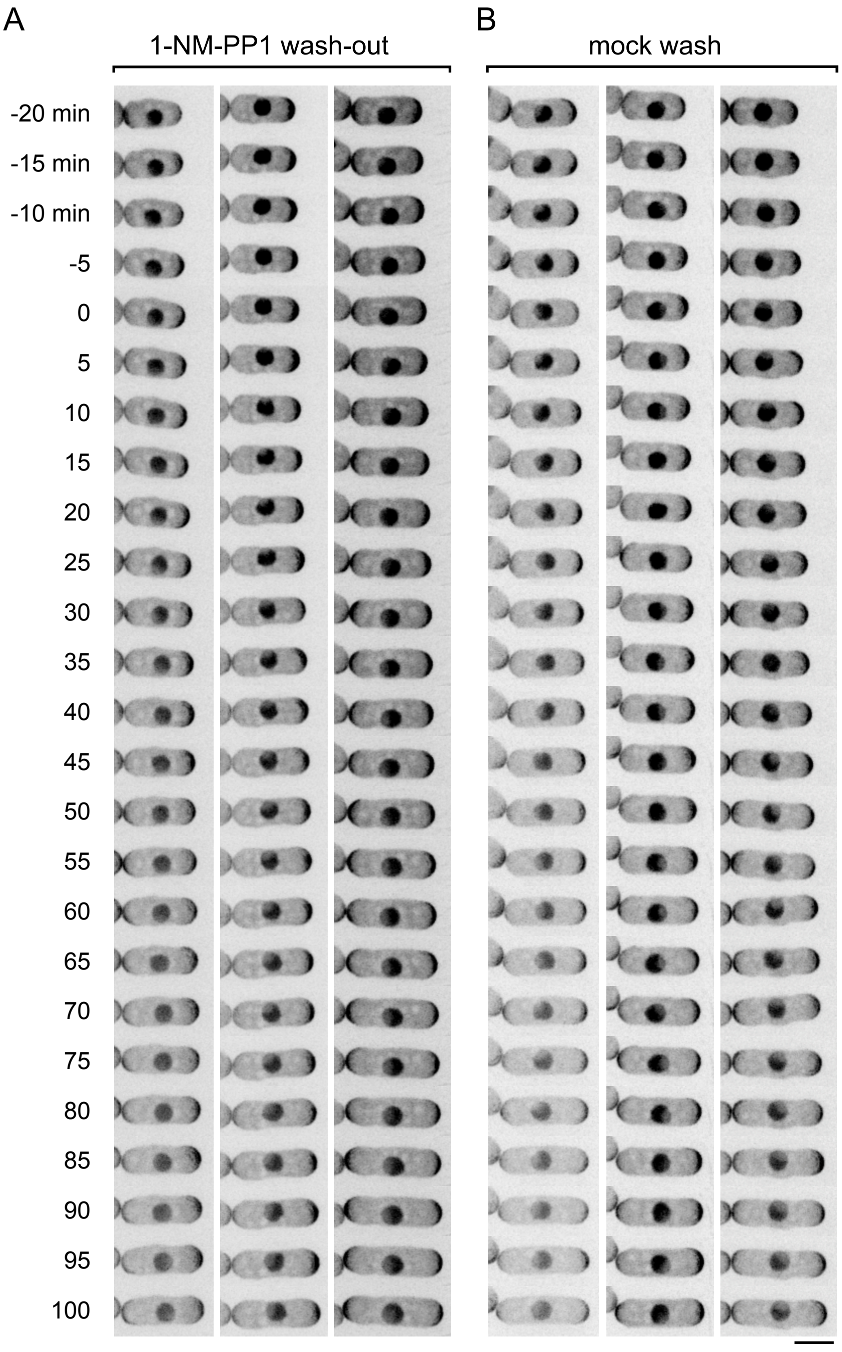


**Extended data Figure S5. Cell polarity in wild-type cells is not altered by washout of AS-kinase inhibitor 1-NM-PP1.**

Timepoints from movies of wild-type (i.e. non-*SISA)* cells expressing Cdc42-GTP reporter CRIB-3mCitrine after (A) wash-out of 2 μM 1-NM-PP1 from growth medium or (B) mock wash into growth medium still containing 2 μM 1-NM-PP1. Times shown are minutes relative to wash-out or mock wash. Because of multiple washes, the zero timepoint was defined as the first timepoint immediately after washing (see Methods). After washout, CRIB-3mCitrine remains at cell tips, and cells continue to elongate. Scale bar, 5 μm.

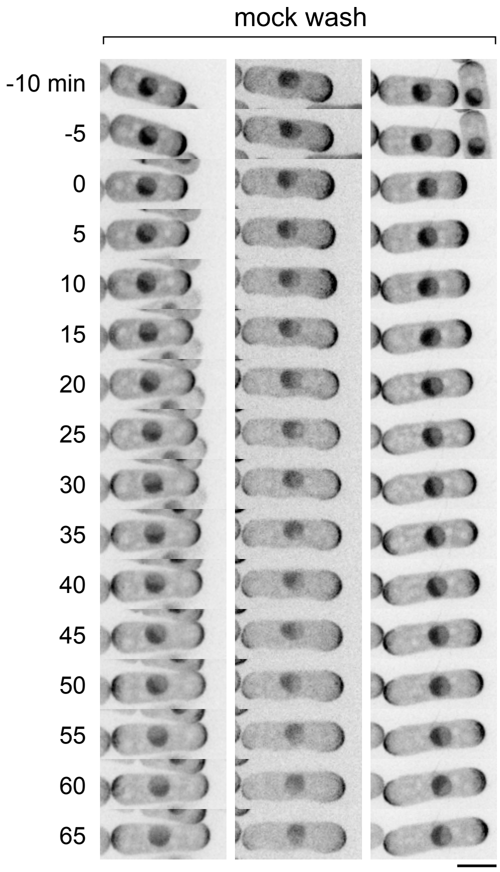


**Extended data Figure S6. No changes in Cdc42-GTP localization after mock wash in *SISA4 atf1*Δ cells.**
Timepoints from movies of *SISA4 atf1*Δ cells expressing Cdc42-GTP reporter CRIB-3mCitrine after mock wash from growth medium containing 2 μM 1-NM-PP1 into growth medium still containing 2 μM 1-NM-PP1. CRIB-3mCitrine remains at cell tips, and cells continue to elongate. Scale bar, 5 μm.

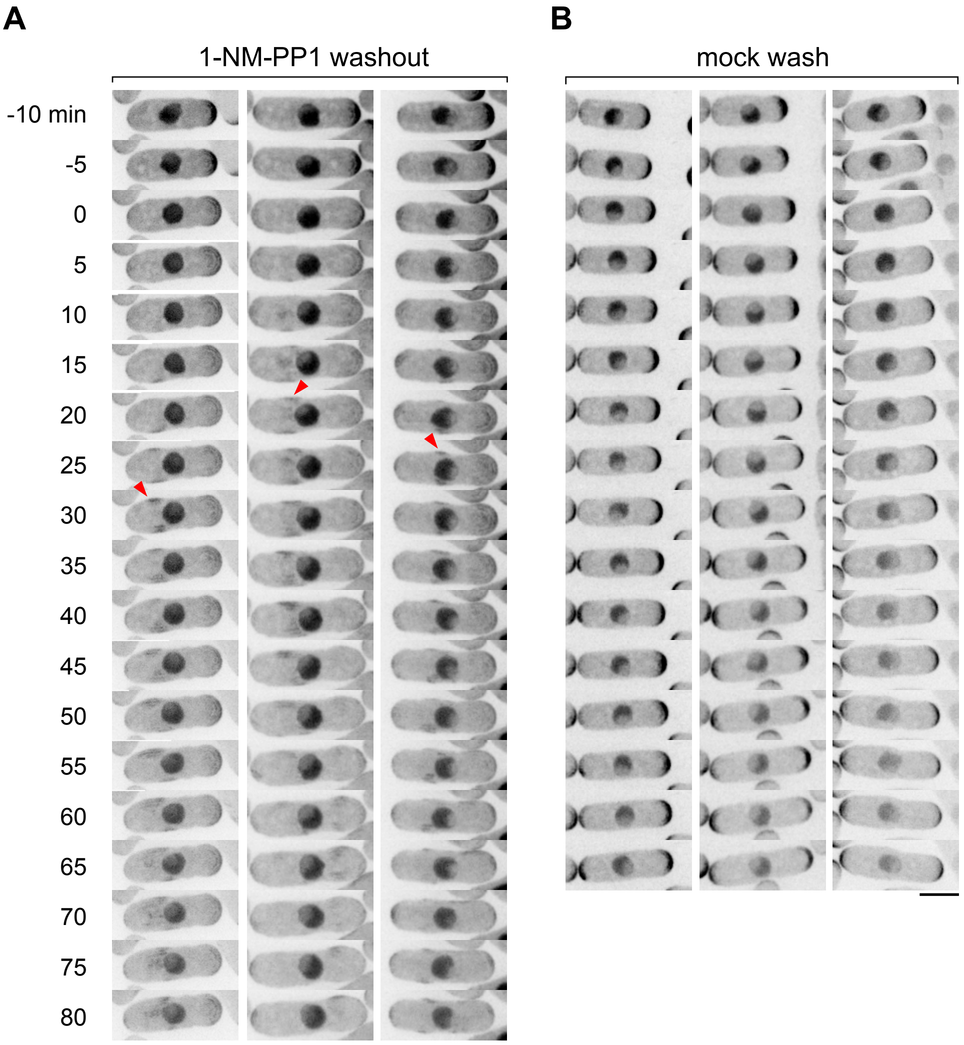


**Extended data Figure S7. Changes in Cdc42-GTP localization at cell tips and cell sides after Sty1 hyperactivation in *SISA4 pcr1*Δ cells.**

Timepoints from movies of *SISA4 pcr1*Δ cells expressing Cdc42-GTP reporter CRIB-3mCitrine upon (A) washout of 2 μM 1-NM-PP1 from growth medium and (B) mock wash into growth medium still containing 2 μM 1-NM-PP1. Note decreased CRIB-3mCitrine at cell tips after washout, appearance of CRIB-3mCitrine patches on cell sides (arrowheads) and cessation of cell elongation. Scale bar, 5 μm.

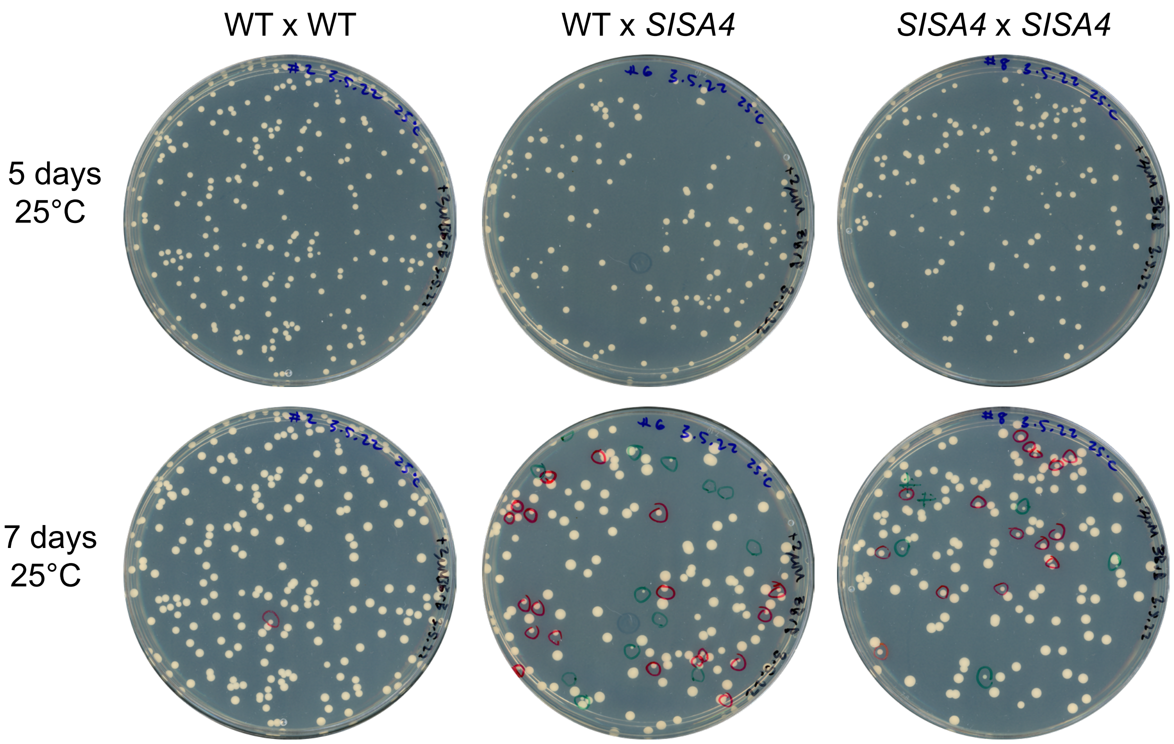


**Extended data Figure S8. Heterogeneous colony size after germination of spores from crosses of wild-type and *SISA4* cells.**

Plate scans of the indicated crosses involving wild-type (WT; non-*SISA*) and *SISA4* strains, after incubation for 5 and 7 days at 25°C on YE5S plates containing 2 µM AS-kinase inhibitor 3-BrB-PP1. Red circles indicate colonies that were smaller than normal after 5 days. Green circles indicate colonies not visible by eye after 5 days but visible after 7 days. In WT x *SISA4* crosses, the majority of SISA4 progeny are found among the slowest-growing colonies (see Table 1), while in *SISA4* x *SISA4* crosses, in which all progeny are *SISA4*, most (but not all) colonies grow at normal rates (see Table 2).

**Extended data Table S1. Tetrad analysis for *SISA4* strain construction**
**(KS11171 *h+ sty1-T97A wis1-DD* x KS11181 *h- sty1-T97A wis1-DD pyp1Δ pyp2Δ*)**

| Col. No. | Col. Pos. | *pyp1* | *pyp2* | *wis1* | Phenotype on YE5S plate |
|---|---|---|---|---|---|
| 1 | AA1 | + | + | DD | alive |
| 2 | AA2 | Δ | Δ | + | alive |
| 3 | AA3 | + | + | + | alive |
| 4 | AA4 | Δ | Δ | DD | dead; normal/swollen |
| 5 | AB1 | Δ | Δ | + | alive |
| 6 | AB2 | + | + | DD | alive |
| - | (AB3) | + | + | + | - |
| 7 | AB4 | Δ | Δ | DD | dead; normal/swollen |
| 8 | AC1 | + | + | DD | alive |
| 9 | AC2 | + | + | DD | alive |
| 10 | AC3 | Δ | Δ | + | alive |
| 11 | AC4 | Δ | Δ | + | alive |
| 12 | AD1 | + | + | + | alive |
| 13 | AD2 | Δ | + | DD | dead; peanut/swollen |
| 14 | AD3 | + | Δ | + | alive |
| 15 | AD4 | Δ | Δ | DD | dead; normal/swollen |
| 16 | AE1 | Δ | Δ | DD | dead; normal/swollen |
| 17 | AE2 | + | + | + | alive |
| 18 | AE3 | + | + | + | alive |
| 19 | AE4 | Δ | Δ | DD | dead; normal/swollen |
| - | (AF1) | + | + | ? | - |
| - | (AF2) | + | + | ? | - |
| 20 | AF3 | Δ | Δ | DD | dead; normal/swollen |
| 21 | AF4 | Δ | Δ | + | alive |
| 22 | AG1 | + | + | DD | alive |
| 23 | AG2 | + | + | + | alive |
| 24 | AG3 | Δ | Δ | + | alive |
| 25 | AG4 | Δ | Δ | DD | dead; normal/swollen |
| - | (AI1) | + | + | DD | - |
| 26 | AI2 | Δ | Δ | DD | dead; normal/swollen |
| 27 | AI3 | + | + | + | alive |
| 28 | AI4 | Δ | Δ | + | alive |
| - | (AA7) | ? | ? | ? | - |
| - | (AA8) | ? | ? | ? | - |
| 29 | AA9 | Δ | Δ | DD | dead; normal/swollen |
| 30 | AA10 | + | + | + | alive |
| 31 | AB7 | + | + | + | alive |
| 32 | AB8 | + | Δ | DD | alive |
| 33 | AB9 | Δ | + | DD | dead; peanut/swollen |
| 34 | AB10 | Δ | Δ | + | alive |
| 35 | AC7 | + | + | + | alive |
| 36 | AC8 | + | + | DD | alive |
| 37 | AC9 | Δ | Δ | + | alive |
| 38 | AC10 | Δ | Δ | DD | dead; normal/swollen |
| 39 | AD7 | Δ | Δ | DD | dead; normal/swollen |
| 40 | AD8 | + | + | + | alive |
| 41 | AD9 | + | + | + | alive |
| 42 | AD10 | Δ | Δ | DD | dead; normal/swollen |
| 43 | AE7 | + | Δ | + | alive |
| 44 | AE8 | Δ | Δ | + | alive |
| 45 | AE9 | Δ | + | DD | dead; peanut/swollen |
| 46 | AE10 | + | + | DD | alive |

| Col. No. | Col. Pos. | *pyp1* | *pyp2* | *wis1* | Phenotype on YE5S plate |
|---|---|---|---|---|---|
| 47 | AF7 | + | + | + | alive |
| 48 | AF8 | Δ | Δ | DD | dead; normal/swollen |
| 49 | AF9 | Δ | Δ | DD | dead; normal/swollen |
| 50 | AF10 | + | + | + | alive |
| 51 | AG7 | + | + | + | alive |
| 52 | AG8 | + | Δ | DD | alive |
| 53 | AG9 | Δ | Δ | + | alive |
| 54 | AG10 | Δ | + | DD | dead; peanut/swollen |
| 55 | AH7 | + | + | + | alive |
| - | (AH8) | Δ | Δ | DD | - |
| 56 | AH9 | Δ | Δ | DD | dead; normal/swollen |
| 57 | AH10 | + | + | + | alive |
| 58 | AI7 | + | + | + | alive |
| - | (AI8) | Δ | Δ | + | - |
| 59 | AI9 | + | + | DD | alive |
| 60 | AI10 | Δ | Δ | DD | dead; normal/swollen |
| 61 | BA1 | Δ | Δ | DD | dead; normal/swollen |
| 62 | BA2 | + | + | + | alive |
| - | (BA3) | + | + | DD | - |
| 63 | BA4 | Δ | Δ | + | alive |
| 64 | BB1 | + | + | DD | alive |
| 65 | BB2 | + | + | DD | alive |
| 66 | BB3 | Δ | Δ | + | alive |
| - | (BB4) | Δ | Δ | + | - |
| 67 | BC1 | + | Δ | + | alive |
| 68 | BC2 | Δ | + | DD | dead; peanut/swollen |
| 69 | BC3 | Δ | Δ | + | alive |
| 70 | BC4 | + | + | DD | alive |
| 71 | BD1 | + | Δ | DD | alive |
| - | (BD2) | Δ | Δ | DD | - |
| 72 | BD3 | Δ | + | + | alive |
| 73 | BD4 | + | + | + | alive |
| 74 | BE1 | Δ | + | + | alive |
| 75 | BE2 | + | + | DD | alive |
| - | (BE3) | + | Δ | DD | - |
| 76 | BE4 | Δ | Δ | + | alive |
| 77 | BF1 | + | + | DD | alive |
| 78 | BF2 | Δ | Δ | + | alive |
| 79 | BF3 | + | + | DD | alive |
| - | (BF4) | Δ | Δ | + | - |
| 80 | BG1 | + | Δ | + | alive |
| 81 | BG2 | Δ | Δ | + | alive |
| 82 | BG3 | Δ | + | DD | dead; peanut/swollen |
| 83 | BG4 | + | + | DD | alive |
| 84 | BH1 | Δ | Δ | DD | dead; normal/swollen |
| 85 | BH2 | + | Δ | DD | alive |
| 86 | BH3 | + | + | + | alive |
| 87 | BH4 | Δ | + | + | alive |
| 88 | BI1 | + | + | DD | alive |
| 89 | BI2 | Δ | + | DD | dead; peanut/swollen |
| 90 | BI3 | + | Δ | + | alive |
| - | (BI4) | Δ | Δ | + | - |

1. Colonies grouped by tetrad. Col. No. indicates colony number. Col. Pos. indicates colony position on YE5S plus 5 µM 3-BrB-PP1 tetrad germination plates (see Extended data Figure S1).
2. Cell phenotypes scored by microscopy after replica-plating colonies from germination plates to YE5S lacking 3-BrB-PP1 and incubating overnight at 32°C.
3. Alleles of *pyp1*, *pyp2* and *wis1-DD* determined by yeast colony PCR.
4. Green text indicates spores that failed to form colonies on germination plates. Inferred genotypes are shown where possible. “?” indicates genotype inference not possible. Among non-germinating spores, no specific genotypes are significantly overrepresented.
5. Red text indicates recombination between *pyp1* and *pyp2* loci. Recombination was always reciprocal.
6. *sty1-T97A wis1-DD pyp1Δ pyp2Δ* genotype is termed *SISA4*. *sty1-T97A wis1-DD pyp1Δ* genotype is termed *SISA3*.

**Extended data Table S2. Strains used in this work**

| Figure | Strain number | Strain genotype | Source |
|---|---|---|---|
| Fig. 1B | AZ107 (KS7671) | *h+ sty1-T97A ura4-D18* | Zuin et al. *EMBO J* 2010, PMID: 20075862 |
| | Shiozaki lab strain 2081 (KS7902) | *h- wis1-DD:12myc::ura4+ leu1-32 ura4-D18* | Shiozaki et al. *Mol Biol Cell* 1998, PMID: 9614178 |
| | KS7818 | *h- sty1-T97A ade6-M210 leu1-32 ura4-D18* | Lab stock; Mutavchiev et al. *Curr Biol* 2016, PMID: 27746023 |
| | KS515 | *h+ ade6-M216 leu1-32 ura4-D18* | Lab stock |
| | KS11179 | *h- sty1-T97A pyp2Δ ade6-M210 leu1-32 ura4-D18* | This study |
| | KS11181 | *h- sty1-T97A pyp1Δ pyp2Δ ade6-M210 leu1-32 ura4-D18* | This study |
| | KS11169 | *h+ sty1-T97A ade6-M210 leu1-32 ura4-D18* | This study |
| | KS9459 | *h- wis1-DD leu1-32 ura4-D18* | This study |
| | KS11171 | *h+ sty1-T97A wis1-DD ade6-M210 leu1-32 ura4-D18* | This study |
| | KS11183 | *h- sty1-T97A wis1-DD pyp1Δ pyp2Δ ade6-M210 leu1-32 ura4-D18 (SISA4)* | This study |
| | KS11184 | *h+ sty1-T97A wis1-DD pyp1Δ pyp2Δ ade6-M210 leu1-32 ura4-D18 (SISA4)* | This study |
| | KS11186 | *h- sty1-T97A wis1-DD pyp1Δ ade6-M210 leu1-32 ura4-D18 (SISA3)* | This study |
| | KS11187 | *h+ sty1-T97A wis1-DD pyp1Δ ade6-M210 leu1-32 ura4-D18 (SISA3)* | This study |
| Fig. 1C | KS11183 | *h- sty1-T97A wis1-DD pyp1Δ pyp2Δ ade6-M210 leu1-32 ura4-D18 (SISA4)* | This study |
| | KS11186 | *h- sty1-T97A wis1-DD pyp1Δ ade6-M210 leu1-32 ura4-D18 (SISA3)* | |
| Fig. 1D | KS515 | *h+ ade6-M216 leu1-32 ura4-D18* | Lab stock |
| | KS516 | *h- ade6-M210 leu1-32 ura4-D18* | Lab stock |
| | KS11183 | *h- sty1-T97A wis1-DD pyp1Δ pyp2Δ ade6-M210 leu1-32 ura4-D18 (SISA4)* | This study |
| | KS11184 | *h+ sty1-T97A wis1-DD pyp1Δ pyp2Δ ade6-M210 leu1-32 ura4-D18 (SISA4)* | This study |
| | KS11186 | *h- sty1-T97A wis1-DD pyp1Δ ade6-M210 leu1-32 ura4-D18 (SISA3)* | This study |
| | KS11187 | *h+ sty1-T97A wis1-DD pyp1Δ ade6-M210 leu1-32 ura4-D18 (SISA3)* | This study |
| Fig. 2B | KS8226 | *h- sty1-T97A wis1-DD:12myc::ura4+ pyp1::ura4+ pyp2::LEU2 adh13-CRIB-3xmECitrine:LEU2 ura4-D18 leu1-32 SISA* | Lab stock; Mutavchiev et al. *Curr Biol* 2016, PMID: 27746023 |
| | KS11051 | *h- sty1-T97A wis1-DD pyp1Δ pyp2Δ adh13-CRIB-3xmECitrine:LEU2 ade6-M210 ura4-D18 leu1-32 (SISA4)* | This study |
| Fig. 3A, B | KS516 | *h- ade6-M210 leu1-32 ura4-D18* | Lab stock |
| | KS11183 | *h- sty1-T97A wis1-DD pyp1Δ pyp2Δ ade6-M210 leu1-32 ura4-D18 (SISA4)* | This study |
| | KS11186 | *h- sty1-T97A wis1-DD pyp1Δ ade6-M210 leu1-32 ura4-D18 (SISA3)* | This study |
| Fig. 4A, B | KS515 | *h+ ade6-M216 leu1-32 ura4-D18* | |
| | KS11184 | *h+ sty1-T97A wis1-DD pyp1Δ pyp2Δ ade6-M210 leu1-32 ura4-D18 (SISA4)* | |
| | KS11014 | *h+ sty1Δ:kanMX ade6-M210 ura4-D18 leu1-32* | |
| Fig. 5A | KS11147 | *h+ lsd90-3xmNeonGreen:hphMX6 sty1-T97A wis1-DD pyp1Δ pyp2Δ ade6-M210 leu1-32 ura4-D18 (SISA4)* | This study |
| Fig. 5B | KS10652 | *h- lsd90-3xmNeonGreen:hphMX6 ade6-M210 leu1-32 ura4-D18* | This study |
| Fig. 6A | KS11147 | *h+ lsd90-3xmNeonGreen:hphMX6 sty1-T97A wis1-DD pyp1Δ pyp2Δ ade6-M210 leu1-32 ura4-D18 (SISA4)* | This study |
| Fig. 6B | KS10652 | *h- lsd90-3xmNeonGreen:hphMX6 ade6-M210 leu1-32 ura4-D18* | This study |
| Fig. 7A, B | KS11051 | *h- sty1-T97A wis1-DD pyp1Δ pyp2Δ adh13-CRIB-3xmECitrine:LEU2 ade6-M210 ura4-D18 leu1-32 (SISA4)* | This study |
| Fig. 8A, B | KS11051 | *h- sty1-T97A wis1-DD pyp1Δ pyp2Δ adh13-CRIB-3xmECitrine:LEU2 ade6-M210 ura4-D18 leu1-32 (SISA4)* | This study |
| Fig. 9 | KS11377 | *h- atf1Δ::kanMX6 sty1-T97A wis1-DD pyp1Δ pyp2Δ Padh13:CRIB-3xmCitrine:LEU2 ade6-M21X leu1-32 ura4-D18 (SISA4)* | This study |

| | | | |
|---|---|---|---|
| Fig. 10A, E, I | KS11051 | *h- sty1-T97A wis1-DD pyp1Δ pyp2Δ adh13-CRIB-3xmECitrine:LEU2 ade6-M210 ura4-D18 leu1-32 (SISA4)* | This study |
| Fig. 10B, F, J | KS11377 | *h- atf1Δ::kanMX6 sty1-T97A wis1-DD pyp1Δ pyp2Δ Padh13:CRIB-3xmCitrine:LEU2 ade6-M21X leu1-32 ura4-D18 (SISA4)* | This study |
| Fig. 10C, G, K | KS11379 | *h- pcr1Δ::bsdMX6 sty1.T97A wis1DD pyp1Δ pyp2Δ adh13-CRIB-3xmECitrine:LEU2 ade6-M21X leu1-32 ura4-D18 (SISA4)* | This study |
| Fig. 10D, H, L | KS12165 | *h- atf1Δ::kanMX6 pcr1Δ::bsdMX6 sty1.T97A wis1DD pyp1Δ pyp2Δ adh13-CRIB-3xmECitrine:LEU2 ade6-M21X leu1-32 ura4-D18 (SISA4)* | This study |
| Fig. 11 | KS11184 | *h+ sty1-T97A wis1-DD pyp1Δ pyp2Δ ade6-M210 leu1-32 ura4-D18 (SISA4)* | This study |
| | KS516 | *h- ade6-M210 leu1-32 ura4-D18* | Lab stock |
| | KS11184 | *h+ sty1-T97A wis1-DD pyp1Δ pyp2Δ ade6-M210 leu1-32 ura4-D18 (SISA4)* | This study |
| | KS7818 | *h- sty1-T97A ade6-M210 leu1-32 ura4-D18* | Lab stock; Mutavchiev et al. *Curr Biol* 2016, PMID: 27746023 |
| | KS11183 | *h- sty1-T97A wis1-DD pyp1Δ pyp2Δ ade6-M210 leu1-32 ura4-D18 (SISA4)* | This study |
| | KS11171 | *h+ sty1-T97A wis1-DD ade6-M210 leu1-32 ura4-D18* | This study |
| Fig. 12A, B | KS7818 | *h- sty1-T97A ade6-M210 leu1-32 ura4-D18* | |
| | KS515 | *h+ ade6-M216 leu1-32 ura4-D18* | |
| | KS11183 | *h- sty1-T97A wis1-DD pyp1Δ pyp2Δ ade6-M210 leu1-32 ura4-D18 (SISA4)* | |
| | KS11184 | *h+ sty1-T97A wis1-DD pyp1Δ pyp2Δ ade6-M210 leu1-32 ura4-D18 (SISA4)* | |
| Table 1 | KS11184 | *h+ sty1-T97A wis1-DD pyp1Δ pyp2Δ ade6-M210 leu1-32 ura4-D18 (SISA4)* | This study |
| | KS516 | *h- ade6-M210 leu1-32 ura4-D18* | Lab stock |
| Table 2 | KS11183 | *h- sty1-T97A wis1-DD pyp1Δ pyp2Δ ade6-M210 leu1-32 ura4-D18 (SISA4)* | This study |
| | KS11184 | *h+ sty1-T97A wis1-DD pyp1Δ pyp2Δ ade6-M210 leu1-32 ura4-D18 (SISA4)* | This study |
| Extended data Fig. S1A | Shiozaki lab strain 2081 (KS7902) | *h- wis1-DD:12myc::ura4+ leu1-32 ura4-D18* | Shiozaki et al. *Mol Biol Cell* 1998, PMID: 9614178 |
| | KS9459 | *h- wis1-DD leu1-32 ura4-D18* | This study |
| Extended data Fig. S1B | KS11171 | *h+ sty1-T97A wis1-DD ade6-M210 leu1-32 ura4-D18* | This study |
| | KS11181 | *h- sty1-T97A pyp1Δ pyp2Δ ade6-M210 leu1-32 ura4-D18* | This study |
| Extended data Fig. S1C | KS515 | *h+ ade6-M216 leu1-32 ura4-D18* | Lab stock |
| | KS516 | *h- ade6-M210 leu1-32 ura4-D18* | Lab stock |
| | KS11183 | *h- sty1-T97A wis1-DD pyp1Δ pyp2Δ ade6-M210 leu1-32 ura4-D18 (SISA4)* | This study |
| | KS11184 | *h+ sty1-T97A wis1-DD pyp1Δ pyp2Δ ade6-M210 leu1-32 ura4-D18 (SISA4)* | This study |
| | KS11186 | *h- sty1-T97A wis1-DD pyp1Δ ade6-M210 leu1-32 ura4-D18 (SISA3)* | This study |
| | KS11187 | *h+ sty1-T97A wis1-DD pyp1Δ ade6-M210 leu1-32 ura4-D18 (SISA3)* | This study |
| Extended data Fig. S2A, B, C | KS515 | *h+ ade6-M216 leu1-32 ura4-D18* | Lab stock |
| | KS516 | *h- ade6-M210 leu1-32 ura4-D18* | Lab stock |
| | KS11183 | *h- sty1-T97A wis1-DD pyp1Δ pyp2Δ ade6-M210 leu1-32 ura4-D18 (SISA4)* | This study |
| | KS11184 | *h+ sty1-T97A wis1-DD pyp1Δ pyp2Δ ade6-M210 leu1-32 ura4-D18 (SISA4)* | This study |
| | KS11186 | *h- sty1-T97A wis1-DD pyp1Δ ade6-M210 leu1-32 ura4-D18 (SISA3)* | This study |
| | KS11187 | *h+ sty1-T97A wis1-DD pyp1Δ ade6-M210 leu1-32 ura4-D18 (SISA3)* | This study |
| Extended data Fig. S3A, B | KS10652 | *h- lsd90-3xmNeonGreen:hphMX6 ade6-M210 leu1-32 ura4-D18* | This study |

| | | | |
|---|---|---|---|
| Extended data Fig. S3C | KS10862 | *h- sty1Δ::natMX6 lsd90-3xmNeonGreen-Ura4:hphMX6 ade6-M210 leu1-32 ura4-D18* | |
| Extended data Fig. S4A | KS10652 | *h- lsd90-3xmNeonGreen:hphMX6 ade6-M210 leu1-32 ura4-D18* | This study |
| Extended data Fig. S4B | KS10862 | *h- sty1Δ::natMX6 lsd90-3xmNeonGreen-Ura4:hphMX6 ade6-M210 leu1-32 ura4-D18* | This study |
| Extended data Fig. S4C | KS11147 | *h+ lsd90-3xmNeonGreen:hphMX6 sty1-T97A wis1-DD pyp1Δ pyp2Δ ade6-M210 leu1-32 ura4-D18 (SISA4)* | This study |
| Extended data Fig. S5A, B | KS7305 | *h- Padh13:CRIB-3xmCitrine:LEU2 ade6-M210 leu1-32 ura4-D18* | Lab stock; Mutavchiev et al. *Curr Biol* 2016, PMID: 27746023 |
| Extended data Fig. S6 | KS11377 | *h- atf1Δ::kanMX6 sty1-T97A wis1-DD pyp1Δ pyp2Δ Padh13:CRIB-3xmCitrine:LEU2 ade6-M21X leu1-32 ura4-D18 (SISA4)* | This study |
| Extended data Fig. S7A, B | KS11379 | *h- pcr1Δ::bsdMX6 sty1-T97A wis1DD pyp1Δ pyp2Δ Padh13:CRIB-3xmECitrine:LEU2 ade6-M21X leu1-32 ura4-D18 (SISA4)* | This study |
| Extended data Fig. S8 | KS515 | *h+ ade6-M216 leu1-32 ura4-D18* | Lab stock |
| | KS516 | *h- ade6-M210 leu1-32 ura4-D18* | Lab stock |
| | KS11184 | *h+ sty1-T97A wis1-DD pyp1Δ pyp2Δ ade6-M210 leu1-32 ura4-D18 (SISA4)* | This study |

### Extended data Table S3. Oligonucleotides used in this work

| General area | Oligo number | Oligo sequence | Purpose | Notes |
|---|---|---|---|---|
| pyp1Δ and pyp2Δ CRISPR yeast strain construction | OKS4824 | CtagaGGTCTCgGACTGACAATCTGGGGCGAGCGCTGTTTcGAGACCcttCC | Top strand for pyp1Δ sgRNA cloning into pLSB-Nat | |
| pyp1Δ and pyp2Δ CRISPR yeast strain construction | OKS4825 | GGaagGGTCTCgAAACAGCGCTCGCCCCAGATTGTCAGTCcGAGACCtctaG | Bottom strand for pyp1Δ sgRNA cloning into pLSB-Nat | |
| pyp1Δ and pyp2Δ CRISPR yeast strain construction | OKS4826 | TACTTTTTTTTTTAATCATCTCTGCTTCTTTTTTAAGGCCAAATATTCTTAATACAAAACATTCATCTAAAAAACCACGAAATTTTTGACTGGATTTTTC | Homology repair template forward primer for pyp1Δ | |
| pyp1Δ and pyp2Δ CRISPR yeast strain construction | OKS4827 | AGACACTTTACAAGTACAAGAAATAAAGGAATCGATTAAAACACGAATATATATTGCCAAGAAAAATCCAGTCAAAAATTTCGTGGTTTTTTAGATGAAT | Homology repair template reverse primer for pyp1Δ | |
| pyp1Δ and pyp2Δ CRISPR yeast strain construction | OKS4830 | CtagaGGTCTCgGACTTGCCTGCCAGGCCCCTAAGCGTTTcGAGACCcttCC | Top strand for pyp2Δ sgRNA cloning into pLSB-Nat | |
| pyp1Δ and pyp2Δ CRISPR yeast strain construction | OKS4831 | GGaagGGTCTCgAAACGCTTAGGGGCCTGGCAGGCAAGTCcGAGACCtctaG | Bottom strand for pyp2Δ sgRNA cloning into pLSB-Nat | |
| pyp1Δ and pyp2Δ CRISPR yeast strain construction | OKS4834 | AGCTTGGCCAATCTTGTCTCTGTCCCTTTTACTTAGGATATAACATCTTCCAAGGTGTTTTCAAAAATTTTGTGCTACTACGAAACGACTGTTCTTTAAT | Homology repair template forward primer for pyp2Δ | |
| pyp1Δ and pyp2Δ CRISPR yeast strain construction | OKS4835 | AAAAAATGAGGAATTACACAATCTCACATAAATAGAACATAGTGTGTACAAACACAGAAAATTAAAGAACAGTCGTTTCGTAGTAGCACAAAATTTTTGA | Homology repair template reverse primer for pyp2Δ | |
| pyp1Δ and pyp2Δ CRISPR yeast strain construction | WC98 | gacggccagtgttgtaaaacg | Forward sequencing primer for sgRNA insert | Torres-Garcia et al., Wellcome Open Res 2020, PMID: 33313420 |
| pyp1Δ and pyp2Δ CRISPR yeast strain construction | WC99 | tacactttatgcttccggctc | Reverse sequencing primer for sgRNA insert | Torres-Garcia et al., Wellcome Open Res 2020, PMID: 33313420 |
| SISA4 genotyping | OKS3898 | GTGGCTGATATATCCAAAGAT | Yeast colony PCR forward primer to confirm wis1-DD | product 739 bp; allele-specific to wis1-DD |
| SISA4 genotyping | OKS3951 | AACTTTCGTTTGCACTTC | Yeast colony PCR reverse primer to confirm wis1-DD | product 739 bp |
| SISA4 genotyping | OKS4856 | CCTCATCCCCTCTAAAACTCTTCC | Yeast colony PCR forward primer to confirm pyp1Δ | product 719 bp |
| SISA4 genotyping | OKS4853 | ACGACCACTTCGCTTAAACTCTAT | Yeast colony PCR reverse primer to confirm pyp1Δ | product 719 bp |
| SISA4 genotyping | OKS4860 | CGTCGCCCAGGAGTTTCTTT | Yeast colony PCR forward primer to confirm pyp2Δ | product 806 bp |
| SISA4 genotyping | OKS4861 | TTGCGTCTCGGCTACGATAA | Yeast colony PCR reverse primer to confirm pyp2Δ | product 806 bp |
| SISA4 genotyping | OKS4971 | GCGATATTTTTATTTCTCCCTTTGAAGATATTTATTTTGTAG | Yeast colony PCR forward primer to confirm sty1-T97A | product 750 bp; allele-specific to sty1-T97A and NOT sty1+ |
| SISA4 genotyping | OKS4987 | CTCCCTTTGAAGATATTTATTTTGTAA | Yeast colony PCR forward primer to confirm sty1+ | product 750 bp; allele-specific to sty1+ and NOT sty1-T97A |
| SISA4 genotyping | OKS4974 | GGATTGCAGTTCATTATCCATGTTGTGAAA | Yeast colony PCR reverse primer to confirm sty1+ and sty1-T97A | product 750 bp; common to sty1+ and sty1-T97A |
| atf1Δ yeast strain construction | OKS2558 | TTCGTTATTTAAGCGTGTGTGATTTTTAACGTCATCTCTTAATACATTTTGTCAAGTGTTAAAGCATAAATTCTTCAATTCGGATCCCCGGGTTAATTAA | Forward PCR primer to make atf1Δ | |
| atf1Δ yeast strain construction | OKS2559 | AGGGAAAGTCAAAGGGAAGTAAAAACGTTATTCTTCATTACAACCATCACAGACTGAGACCTTTTCAGATCAAAAACAGTGAATTCGAGCTCGTTTAAAC | Reverse PCR primer to make atf1Δ | |
| atf1Δ yeast strain construction | OKS2560 | GGAAAGCCATTAATTTCTTGTAGTTGTT | Yeast colony PCR forward primer to confirm atf1Δ | |
| atf1Δ yeast strain construction | OKS2561 | GGGAAGAAATATAAAAAGAGAAGCCTAA | Yeast colony PCR reverse primer to confirm atf1Δ | |
| pcr1Δ yeast strain construction | OKS4989 | CATCTCCCCCTTTGTTTCCTTTGTTATATTTTTGTTTGTGATCATCTGATTCCCCCTTTTCTATACATTGATTATTTAAGCGGATCCCCGGGTTAATTAA | Forward PCR primer to make pcr1Δ | |
| pcr1Δ yeast strain construction | OKS4990 | AGCCATAAATCTACATGCAACATCACATTAAAATCAATTGCTTTATTGTAAAGGGGGGTAATGAAAACAAATGTTCTTTTGAATTCGAGCTCGTTTAAAC | Reverse PCR primer to make pcr1Δ | |
| pcr1Δ yeast strain construction | OKS2940 | AGTAATATCATGCGTCAATCGTATG | Yeast colony PCR forward primer to confirm pcr1Δ | |
| pcr1Δ yeast strain construction | OKS4946 | AATCTGTACATTTAGAAAGAAGAAC | Yeast colony PCR reverse primer to confirm pcr1Δ | |
| Lsd90-3xmNeonGreen strain construction | OKS4544 | GCAAGGAACGTAGAAGCAGCACCAGCAGCGGTCATGGCTTGATGAACAATGTTCGTCATGCCTTGGGTATGTCCAACAAGCGGATCCCCGGGTTAAT | Forward PCR primer to make Lsd90-3xmNeonGreen | |
| Lsd90-3xmNeonGreen strain construction | OKS4545 | CAATAAAAACCTTTTTTCATAACGGATTCACGAGATAATGAAGGTTACGTCCAAAAATTTTGATATTGCAAGTATTCCCTGAATTCGAGCTCGTTTAAAC | Reverse PCR primer to make Lsd90-3xmNeonGreen | |
